# Ca^2+^-sensing receptor regulates neuronal excitability via Kv7 channel and G_i/o_ protein signalling

**DOI:** 10.1101/2023.07.07.548061

**Authors:** Nontawat Chuinsiri, Nannapat Siraboriphantakul, Luke Kendall, Polina Yarova, Christopher J. Nile, Bing Song, Ilona Obara, Justin Durham, Vsevolod Telezhkin

## Abstract

Neuropathic pain, a debilitating condition with unmet medical needs, can be charactarised as hyperexcitability of nociceptive neurons caused by dysfunction of ion channels. Voltage-gated potassium channel type 7 (Kv7), responsible for maintaining neuronal resting membrane potential and thus neuronal exitability, resides under tight control of G protein-coupled receptors (GPCR). Calcium-sensing receptor (CaSR) is a GPCR that is known to regulate activity of numerous ion channels, but whether CaSR could control Kv7 channel function has been unexplored until now. Our results demonstrate that CaSR is expressed in recombinant cell models, human induced pluripotent stem cell (hiPSC)-derived nociceptive-like neurons and mouse dorsal root ganglia neurons, and its activation induced depolarisation via Kv7.2/7.3 channel inhibition. The CaSR-Kv7.2/7.3 channel crosslink was mediated via the G_i/o_ protein/adenylate cyclase/cyclic adenosine monophosphate/protein kinase A signalling cascade. Suppression of CaSR function rescued hiPSC-derived nociceptive-like neurons from algogenic cocktail-induced hyperexcitability. To conclude, this study demonstrates that CaSR-Kv7.2/7.3 channel crosslink via the G_i/o_ protein signalling pathway effectively regulates neuronal excitability, providing a feasible pharmacological target for neuronal hyperexcitability management in neuropathic pain.

## 1 Introduction

Neuropathic pain, defined as a pain caused by a lesion or disease affecting the somatosensory system, exerts significant impacts on individuals’ quality of life (Oberoi *et al*, 2014; Shueb *et al*, 2015) and has serious socio-economic burden (Schaefer *et al*, 2014). The pathophysiological mechanisms of neuropathic pain are poorly understood, but existing evidence points towards hyperexcitability of nociceptive neurons associated with functional and structural alterations of neuronal ion channels such as voltage-gated sodium (Nav), voltage-gated potassium (Kv), and voltage-gated calcium (Cav) channels (Iwata *et al*, 2011; Li *et al*, 2014; Siqueira *et al*, 2009; Takeda *et al*, 2011). Carbamazepine, a Nav channel blocker, and gabapentinoids, a Cav channel blocker, have been used for neuropathic pain management. However, these medications are not always effective in producing a satisfactory pain reduction and often are associated with unwanted side effects (Haviv *et al*, 2014), suggesting that direct targeting of specific ion channels may not be an optimal solution. Upstream regulatory mechanisms of the key ion channels via G protein-coupled receptors (GPCRs) may be implicated in the pathophysiology of neuronal hyperresponsiveness, thus it could be suggested that GPCRs targeting might provide a more appropriate therapeutic avenue for neuropathic pain management (Pan *et al*, 2008).

Canonical neuronal Kv7 channels (Kv7.2 and Kv7.3 heterotetramers), encoded by the *KCNQ2* and *KCNQ3* genes, specialise in maintaining the resting membrane potential (RMP) and regulating action potential (AP) firing (Peng *et al*, 2017; Tsantoulas & McMahon, 2014). Dysfunction of Kv7 channels underpins several neuronal hyperexcitability disorders, including epileptic seizures and neuropathic pain (Charlier *et al*, 1998; Jones *et al*, 2021; Ling *et al*, 2017; Nardello *et al*, 2020; Schroeder *et al*, 1998). The current passing through Kv7 channels is known as M current (*I*_m_), which is mostly conducted through Kv7.2/7.3 channels in the neurons (Wang *et al*, 1998). The *I*_m_ is characterised by a continuous voltage-dependent outward current activating at subthreshold potentials, approximately –60 mV, and has slow activation-deactivation kinetics (Brown & Adams, 1980; Selyanko *et al*, 1992; Wang *et al*., 1998). Several studies have revealed that Kv7 channels can also be modulated by GPCRs (Greene & Hoshi, 2017). Muscarinic cholinergic M1 receptor activation was first demonstrated to mediate Kv7 channel inhibition via the G_q/11_ protein signalling pathway (Brown & Adams, 1980; Winks *et al*, 2005). Later, activation of bradykinin B2 receptors and purinergic metabotropic P2Y receptors were also shown to induce *I*_m_ inhibition via the G_q/11_ protein (Cruzblanca *et al*, 1998; Gamper & Shapiro, 2003; Liu *et al*, 2010; Zaika *et al*, 2007).

The Ca^2+^-sensing receptor (CaSR) is a GPCR which main physiological function is maintaining the homeostasis of extracellular Ca^2+^ concentration ([Ca^2+^]_o_) (Leach *et al*, 2020). The canonical signalling pathways of CaSR include G_q/11_ protein-dependent increase in intracellular Ca^2+^ concentration ([Ca^2+^]_i_) (Brown *et al*, 1993) and G_i/o_ protein-dependent reduction in 3′,5′-cyclic adenosine monophosphate (cAMP) levels (Chang *et al*, 1998). Since its original discovery, it has become evident that the CaSR is abundantly expressed throughout the nervous system (Brown & Passmore, 2009; Heyeraas *et al*, 2008; Ruat *et al*, 1995), where it regulates synaptic transmission and neuronal activity (Jones & Smith, 2016), whilst its dysfunction has been linked to seizures (Kapoor *et al*, 2008). The CaSR can regulate neuronal excitability via several ion channels, *e.g.* non-selective cation channels and Ca^2+^-activated K^+^ channels (Jones & Smith, 2016; Vassilev *et al*, 1997; Ye *et al*, 1997). Negative allosteric modulators (NAMs) of the CaSR, more commonly known as calcilytics, have demonstrated promising therapeutic uses in certain neurological diseases, *e.g.* traumatic brain injury and Alzheimer’s disease (Armato *et al*, 2013; Bai *et al*, 2015; Kim *et al*, 2013; Wang *et al*, 2020; Xue *et al*, 2017), whilst positive allosteric modulators (PAMs) or calcimimetics were suggested as neuroblastoma therapy (Rodríguez-Hernández *et al*, 2016).

To the best of our knowledge, no direct investigation of a role that CaSR may play in neuropathic pain development has yet been undertaken. However, the well-established interplay between the CaSR and pro-inflammatory cytokines (Iamartino & Brandi, 2022) and the significance of the inflammatory process underlying many neuropathic pain conditions (Ellis & Bennett, 2013; Li *et al*, 2023) suggests the CaSR as a potential candidate for neuropathic pain therapy. Moreover, based on the evidence that neuronal hyperexcitability is associated with dysfunctions of the CaSR (Kapoor *et al*., 2008) and neuronal Kv7.2/7.3 channels (Chokvithaya *et al*, 2023; Maghera *et al*, 2020; Schroeder *et al*., 1998), it is conceivable that a crosslink between the CaSR and Kv7.2/7.3 channels may exist and contribute to the regulation of neuronal excitability. This study aimed to establish the molecular pathway linking the CaSR and Kv7.2/7.3 channels, and explore the potential of calcilytics in suppressing neuronal hyperexcitability using a human induced pluripotent stem cell (hiPSC)-derived nociceptive-like neuronal model.

## 2 Results

### 2.1 Activation of the CaSR attenuates the *I*_m_ and causes depolarisation through neuronal Kv7.2/7.3 channels

The functional crosslink between the CaSR and Kv7.2/7.3 channels was investigated using several cellular models: Chinese hamster ovary cells stably expressing Kv7.2/7.3 channels (CHO-Kv7.2/7.3), human embryonic kidney 293 cells stably expressing the CaSR (HEK-CaSR), mouse dorsal root ganglia (DRG) neurons and hiPSC-derived nociceptive-like neurons. Molecular expression of the CaSR in CHO-Kv7.2/7.3 was verified with immunocytochemistry (Fig 1A); HEK-CaSR were used as a positive control (Fig 1B).

**Figure 1.**
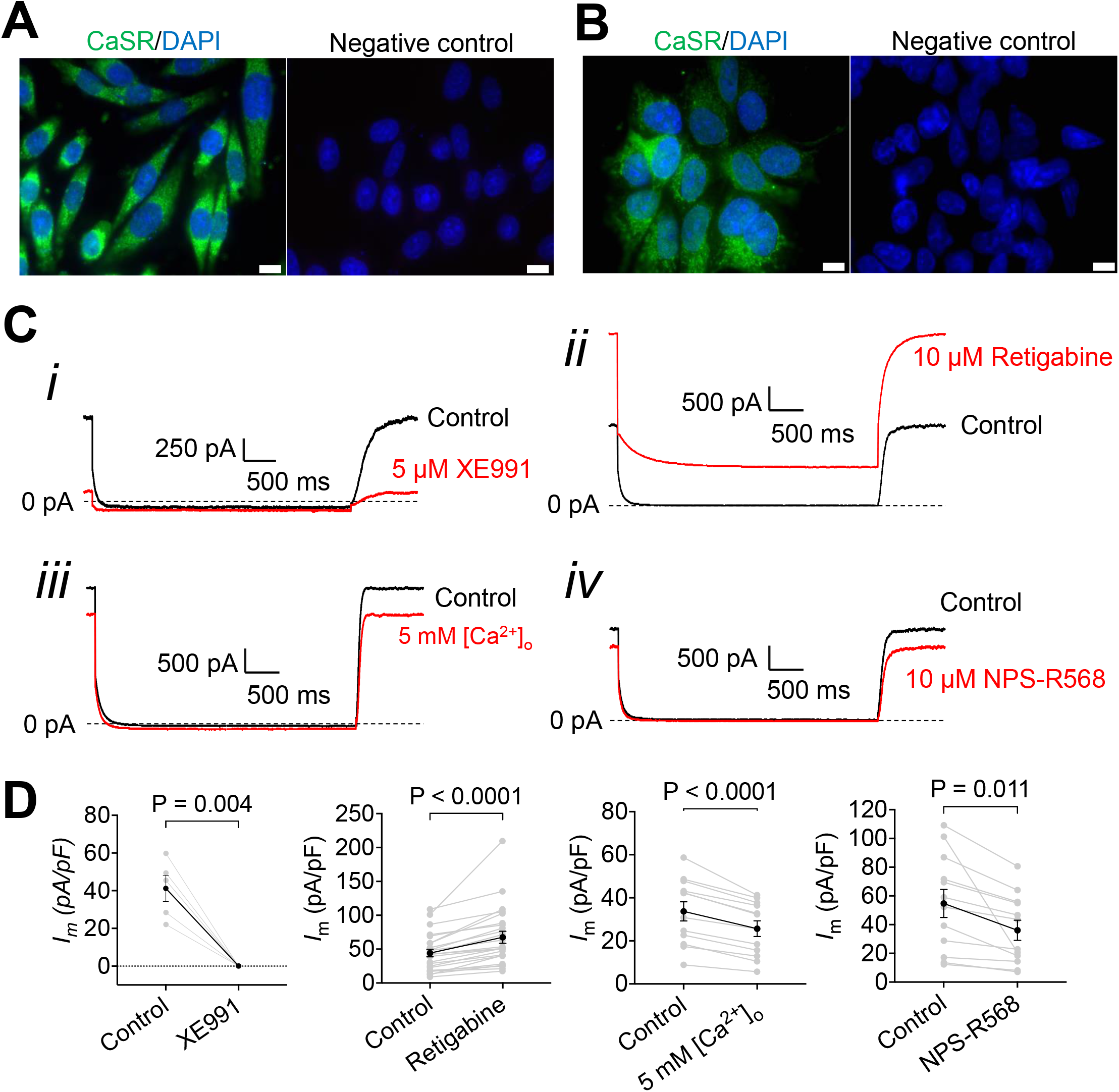
Molecular expression of the CaSR and its effect on the *I*_m_. A and B Immunofluorescence images show the CaSR (ab19347; green) expressed in CHO-Kv7.2/7.3 (A) and HEK-CaSR (B). Images were obtained with Olympus BX61 fluorescence microscope and are representative of three independent experiments. Scale bars: 10 µm. C Exemplar traces show the *I*_m_ of CHO-Kv7.2/7.3 in control (0.5 mM [Ca^2+^]_o_) and in the presence of 5 μM XE991 (i), 10 μM retigabine (ii), 5 mM [Ca^2+^]_o_ (iii) and 10 μM NPS-R568 (iv). D Graphs compare the *I*_m_ of CHO-Kv7.2/7.3 in control (0.5 mM [Ca^2+^]_o_) and in the presence of 5 μM XE991 (n=5 cells), 10 μM retigabine (n=24 cells), 5 mM [Ca^2+^]_o_ (n=12 cells) and 10 μM NPS-R568 (n=12 cells). Data information: In D, data are shown as means±SEM. Statistical analyses were performed using two-tailed paired t-test (for XE991, 5 mM [Ca^2+^]_o_ and NPS-R568) and Wilcoxon matched-pairs signed rank test (for retigabine).

Using the voltage-stepped deactivation protocol, the effects of high [Ca^2+^]_o_ (5 mM) and NPS-R568, a PAM of the CaSR, on the *I*_m_ of CHO-Kv7.2/7.3 were examined in the whole-cell configuration mode. CHO-Kv7.2/7.3 exhibited an outward *I*_m_ of 43.7±4.8 pA/pF (n=29 cells) when bathed in a standard solution (0.5 mM [Ca^2+^]_o_). Application of 5 µM XE991, a Kv7 channel blocker, and 10 µM retigabine, a Kv7 channel opener, significantly decreased (– 41.1±6.9 pA/pF) and increased (23.2±4.4 pA/pF) the *I*_m_ of CHO-Kv7.2/7.3, respectively. In a similar fashion to XE991, high [Ca^2+^]_o_ and 10 µM NPS-R568 significantly reduced the *I*_m_ of CHO-Kv7.2/7.3 by –8.0±1.1 pA/pF and –18.7±6.1 pA/pF, respectively (Fig 1C and D).

The effect of the CaSR on the RMP of CHO-Kv7.2/7.3 was investigated in the current-clamp mode. Administration of [Ca^2+^]_o_ (0.1 to 10 mM) caused depolarisation in a concentration-dependent manner with a half maximal effective concentration (EC_50_) of 4.3 mM (Fig 2A and E). The relationship between [Ca^2+^]_o_ and voltage change was then investigated in the presence of NPS-R568 at two concentrations (1 and 10 μM) (Fig 2B and C); the concentration-dependent effect of [Ca^2+^]_o_ on depolarisation was preserved in the presence of NPS-R568 (Fig 2E). Administration of 1 μM NPS-2143, a NAM or calcilytic, caused a hyperpolarising shift of the [Ca^2+^]_o_-voltage change response (Fig 2D and E). The degree of depolarisation by 0.5, 5 and 10 mM [Ca^2+^]_o_ in the presence of NPS-2143 was significantly decreased, compared to the control (Fig 2E).

**Figure 2.**
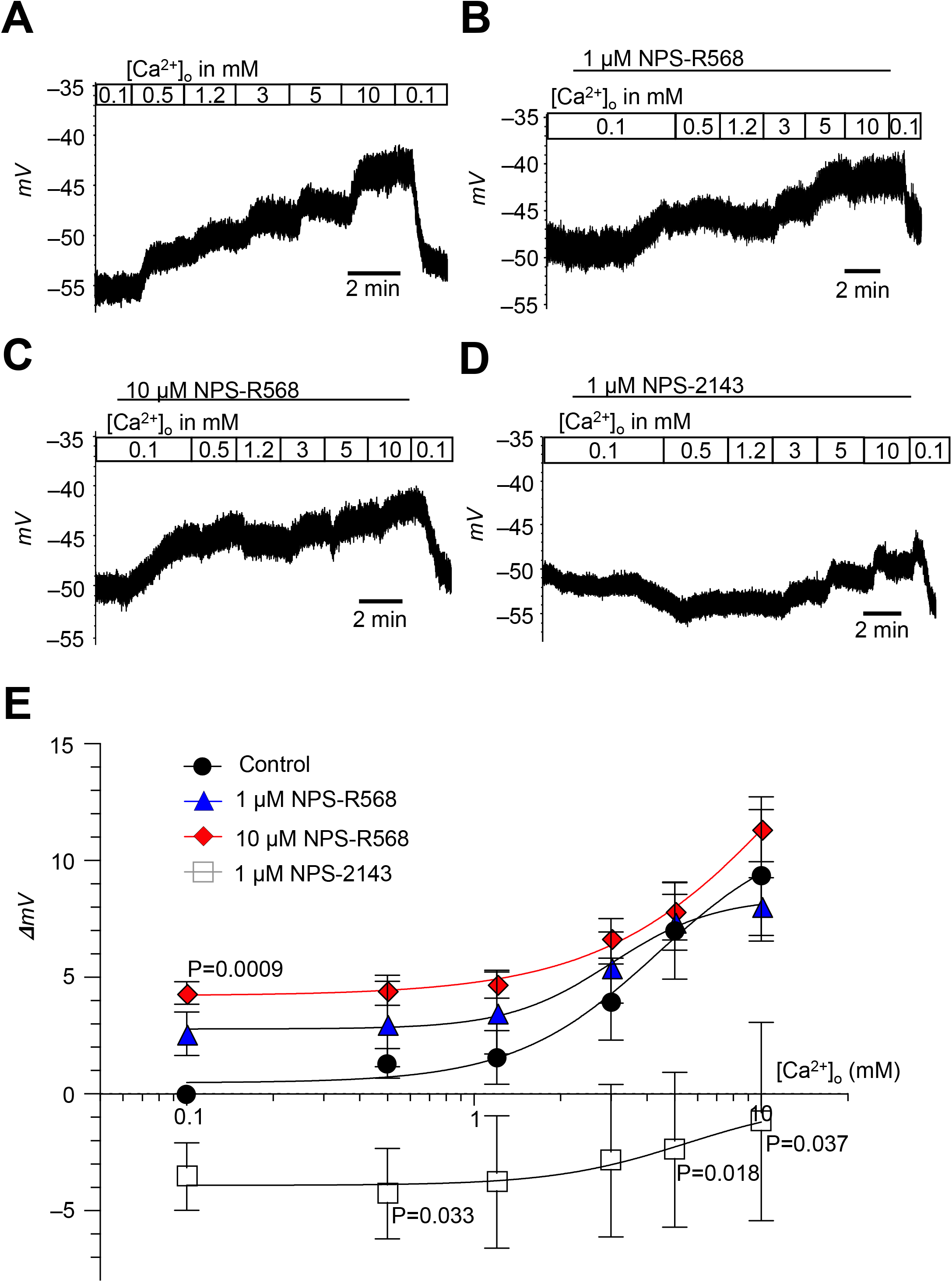
Effect of CaSR modulation on the RMP. A-D Exemplar traces show the effects of [Ca^2+^]_o_ (0.1, 0.5, 1.2, 3, 5 and 10 mM) on the RMP of CHO-Kv7.2/7.3 in the control condition (A), 1 μM NPS-R568 (B), 10 μM NPS-R568 (C), and 1 μM NPS-2143 (D). E Fitted concentration-response curves in each condition are shown (n=5 cells per condition). Data information: Data are shown as means±SEM. In E, one-sample t-test to a theoretical mean of 0 was used to test the effects of 1 μM NPS-R568, 10 μM NPS-R568 and 1 μM NPS-2143 on the RMP in the presence of 0.1 mM [Ca^2+^]_o_, and one-way ANOVA with Dunnett’s multiple comparison test was used to compare the effects of 1 μM NPS-R568, 10 μM NPS-R568 and 1 μM NPS-2143 on the RMP to the control in the presence of 0.5 – 10 mM [Ca^2+^]_o_. Where P is not provided, no significant difference was observed.

To test whether the observed effect of the CaSR on the RMP was mediated through Kv7.2/7.3 channels, transient transfection of HEK-CaSR with a *KCNQ2/3* cDNA plasmid was performed, and the depolarising effect of CaSR activation was investigated. In sham-transfected HEK-CaSR, 10 µM retigabine and 10 µM NPS-R568 did not induce a significant change in the RMP, while depolarisation by 5 mM [Ca^2+^]_o_ (3.8 ±0.7 mV) was observed. In *KCNQ2/3* cDNA-transfected cells, retigabine induced significant hyperpolarisation (–21.9 ±1.4 mV), confirming the functional expression of Kv7.2/7.3 channels. The magnitude of depolarisation by 5 mM [Ca^2+^]_o_ and 10 µM NPS-R568 was also significantly increased by 2.6 ±1.0 mV and 2.1 ±0.9 mV, respectively (Fig 3A and B). These data suggest that CaSR-mediated depolarisation is influenced by the expression of Kv7.2/7.3 channels.

**Figure 3.**
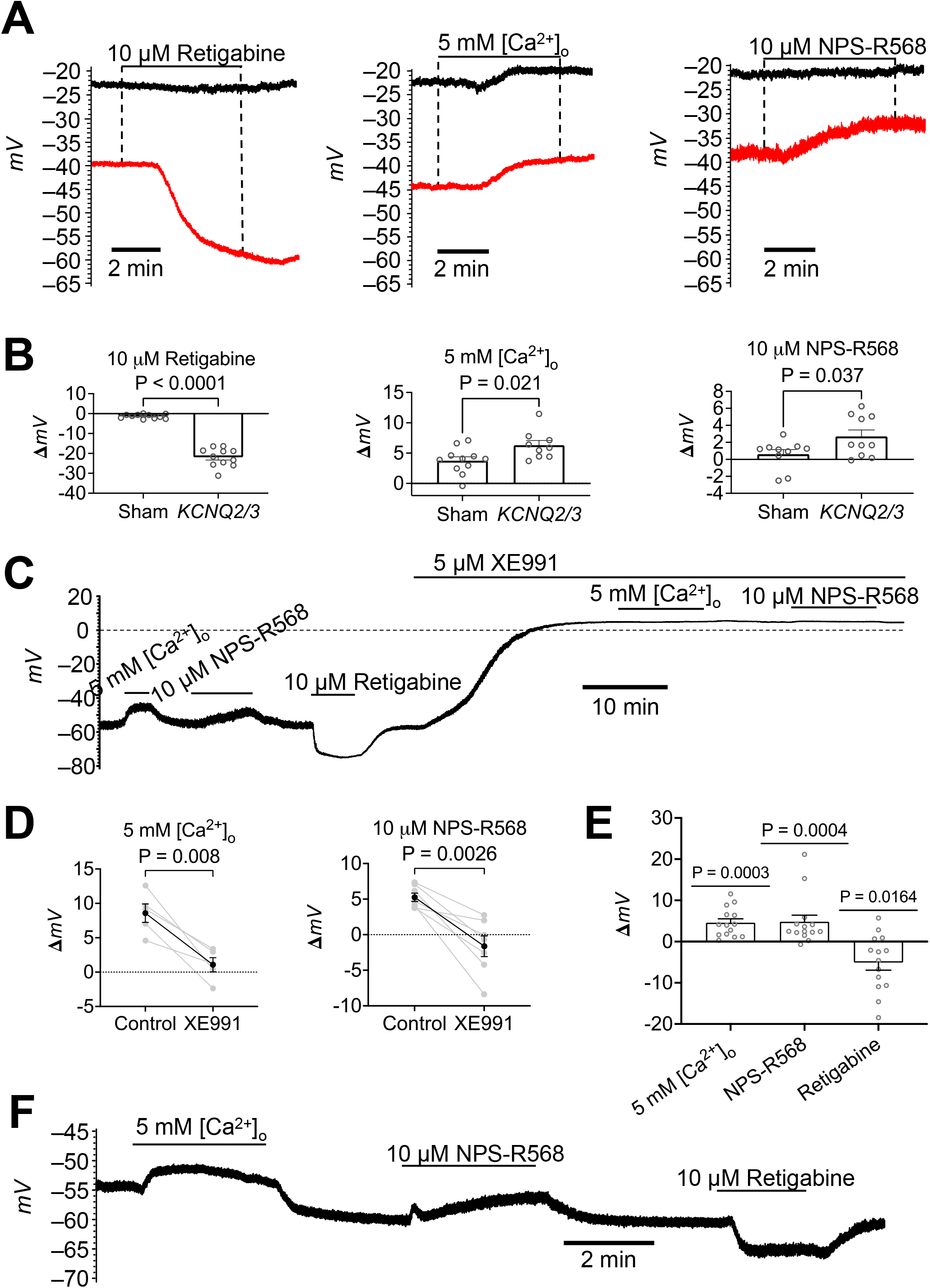
Molecular and pharmacological identification of the functional crosslink between the CaSR and Kv7.2/7.3 channels. A Exemplar traces show the effects of 10 μM retigabine, 5 mM [Ca^2+^]_o_ and 10 μM NPS-R568 on the RMP of sham-transfected HEK-CaSR (black traces) and HEK-CaSR transiently transfected with *KCNQ2/3* cDNA (red traces). B Graphs compare the magnitude of hyperpolarisation by 10 μM retigabine (n=11 cells) and depolarisation by 5 mM [Ca^2+^]_o_ (n=9-11 cells) and 10 μM NPS-R568 (n=10 cells) between sham-transfected and *KCNQ2/3-*transfected HEK-CaSR. C Exemplar trace shows the effects of 5 mM [Ca^2+^]_o_ and 10 μM NPS-R568 on the RMP of CHO-Kv7.2/7.3 in the control condition and in the presence of 5 μM XE991. D Graphs compare the magnitude of depolarisation by 5 mM [Ca^2+^]_o_ (n=5 cells) and 10 μM NPS-R568 (n=7 cells) before and after the application of 5 µM XE991 in CHO-Kv7.2/7.3. E Graph shows the magnitude of depolarisation by 5 mM [Ca^2+^]_o_ and 10 μM NPS-R568 and hyperpolarisation by 10 μM retigabine in mouse DRG neurons (n=14 cells from six mice). F Exemplar trace shows the effects of 5 mM [Ca^2+^]_o_, 10 μM NPS-R568 and 10 μM retigabine on the RMP of a mouse DRG neuron. Data information: Data are shown as means±SEM. Statistical analyses were performed using two-tailed unpaired t-test (B) and two-tailed paired t-test (D). In E, one-sample t-test to a theoretical mean of 0 was used for testing the effects of 5 mM [Ca^2+^]_o_ and retigabine, and the Wilcoxon signed rank test to a theoretical mean of 0 was used for testing the effect of 10 μM NPS-R568.

Further evidence to support the CaSR-Kv7.2/7.3 channel crosslink was derived from experiments comparing the magnitude of depolarisation induced by CaSR activation before and after the application of 5 µM XE991. Increasing [Ca^2+^]_o_ from 0.5 to 5 mM and application of 10 µM NPS-R568 in the presence of 0.5 mM [Ca^2+^]_o_ caused depolarisation in CHO-Kv7.2/7.3. Application of 5 µM XE991 depolarised the cells and significantly attenuated the depolarising effects of 5 mM [Ca^2+^]_o_ by –7.5 ±1.5 mV and NPS-R568 by –6.9 ±1.4 mV (Fig 3C and D).

The CaSR-Kv7 channel crosslink was explored in DRG neurons isolated from six naïve mice. To evaluate the functional expression of the CaSR and Kv7 channels, the effects of 5 mM [Ca^2+^]_o_, 10 μM NPS-R568 and 10 μM retigabine on the RMP were recorded. Overall, application of 5 mM [Ca^2+^]_o_ and NPS-R568 depolarised the neurons by 4.6 ±0.9 mV and 4.8 ±1.6 mV, respectively, whereas retigabine hyperpolarised the neurons by –5.1 ±1.9 mV (Fig 3E and F). Based on the magnitude of voltage change induced by NPS-R568 and retigabine, the neurons were categorised as responding to: NPS-R568 (voltage change > 1 mV), retigabine (voltage change < –1 mV), or both. Of all 14 neurons, eight neurons (57.1%) responded to both NPS-R568 and retigabine, four neurons (28.6%) responded to NPS-R568, and two neurons (14.3%) responded to retigabine. The effect of CaSR activation on the *I*_m_ and RMP was then investigated in nociceptive-like neurons, which were differentiated from HD33n1 hiPSCs and characterised by immunocytochemistry and patch-clamp electrophysiology. Immunocytochemistry confirmed the expression of the CaSR, a neuronal marker microtubule-associated protein 2 (MAP2), a nociceptive marker transient receptor potential vanilloid 1 (TRPV1) (Fig 4A and B) and Kv7.2/7.3 channels (Appendix Fig S1A). Patch-clamp electrophysiology demonstrated the

**Figure 4.**
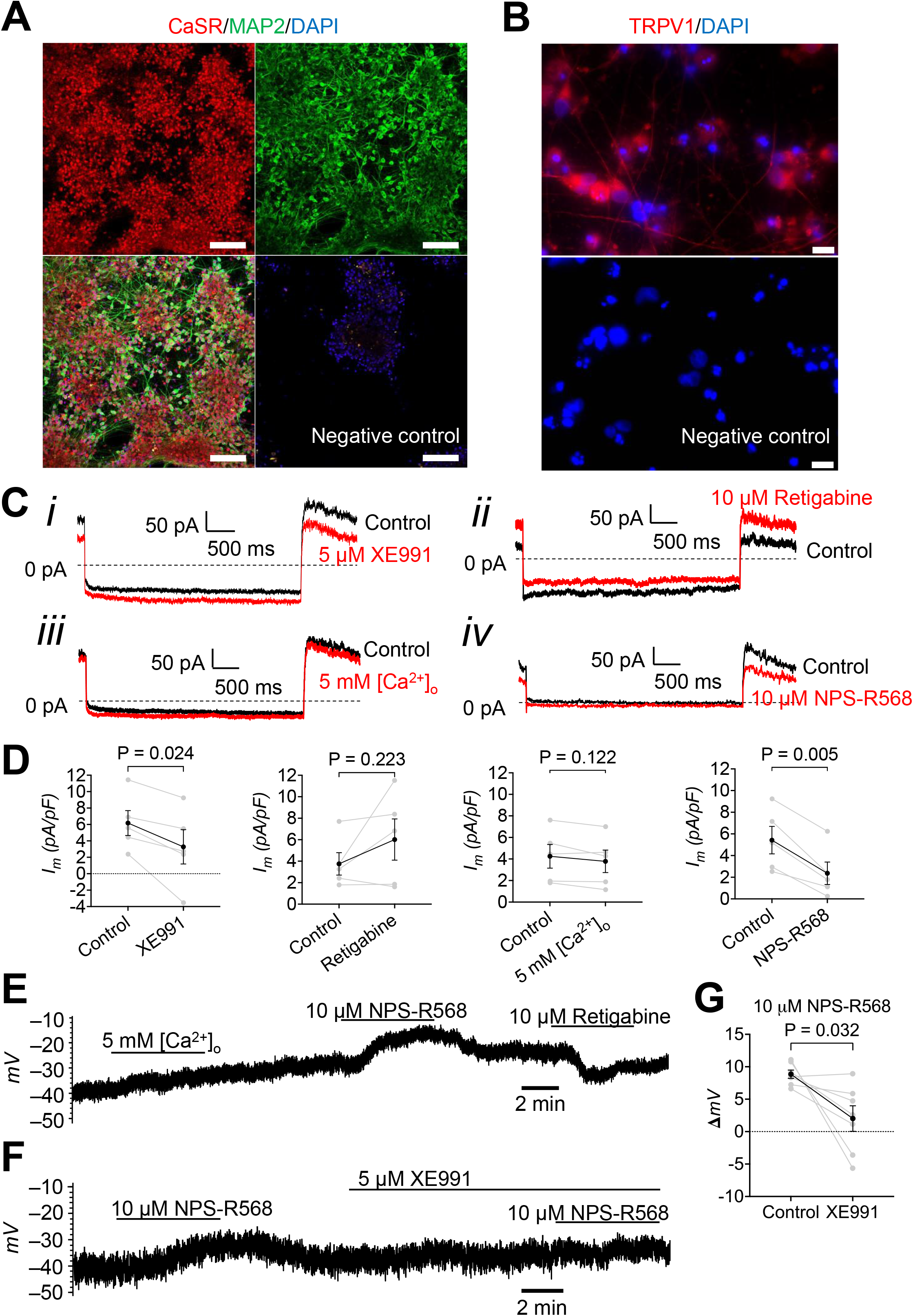
Characterisation of hiPSC-derived neurons and verification of the CaSR-Kv7 channel crosslink. A Immunofluorescence images show the CaSR (SAB4503369; red) and MAP2 (M1406; green) expressed in hiPSC-derived neurons. Images were obtained with Zeiss LSM800 laser confocal scanning microscope and are representative of three independent experiments. Scale bars: 100 µm. B Immunofluorescence image shows TRPV1 (ab3487; red) expressed in hiPSC-derived neurons. Images were obtained with Olympus BX61 fluorescence microscope and are representative of three independent experiments. Scale bars: 10 µm. C Exemplar traces show the *I*_m_ of hiPSC-derived nociceptive-like neurons in control (0.5 mM [Ca^2+^]_o_) and in the presence of 5 μM XE991 (i), 10 μM retigabine (ii), 5 mM [Ca^2+^]_o_ (iii) and 10 μM NPS-R568 (iv). D Graphs compare the *I*_m_ of hiPSC-derived nociceptive-like neurons in control (0.5 mM [Ca^2+^]_o_) and in the presence of 5 μM XE991, 10 μM retigabine, 5 mM [Ca^2+^]_o_ and 10 μM NPS-R568 (n=5 cells). E Exemplar trace shows the effects of 5 mM [Ca^2+^]_o_, 10 μM NPS-R568 and 10 μM retigabine on the RMP of a hiPSC-derived nociceptive-like neurons. F Exemplar trace shows the effect of 10 μM NPS-R568 on the RMP of a hiPSC-derived nociceptive-like neurons before and after the application of 5 μM XE991. G Graph compares the magnitude of depolarisation by 10 μM NPS-R568 before and after the application of 5 µM XE991 in hiPSC-derived nociceptive-like neurons (n=7 cells). Data: information: Data are shown as means±SEM. In D and G, statistical analyses were performed using two-tailed paired t-test.

Nav current, Kv current and Nav channel activation-inactivation profile of the hiPSC-derived nociceptive-like neurons, which were qualitatively similar to those of mouse DRG neurons (Appendix Fig S1B-D). In a standard solution, hiPSC-derived nociceptive-like neurons had the outward *I*_m_ of 5.0±0.7 pA/pF (n=16 cells). Application of 5 µM XE991 significantly reduced the *I*_m_ by –2.9 ±0.8 pA/pF, whereas 10 µM retigabine insignificantly increased the *I*_m_ by 2.3 ±1.6 pA/pF. No apparent effect of 5 mM [Ca^2+^]_o_ on the *I*_m_ was observed. Activation of CaSR with 10 µM NPS-R568 significantly reduced the *I*_m_ by –3.1 ±0.5 pA/pF (Fig 4C and D). In the current-clamp mode, the effect of 5 mM [Ca^2+^]_o_ on the RMP varied between cells, whereas a consistent depolarising effect of 10 µM NPS-R568 was observed (Fig 3E). In the presence of 5 µM XE991, however, NPS-R568-induced depolarisation was significantly attenuated by –6.8 ±2.5 mV in hiPSC-derived nociceptive-like neurons (Fig 4F and G).

### 2.2 CaSR-mediated depolarisation of CHO-Kv7.2/7.3 is sensitive to 8-Br-cAMP but not to NiCl_2_, BAPTA-AM and U73122

To explore the potential molecular pathways linking the CaSR and Kv7.2/7.3 channels, compounds targeting certain canonical G protein signalling pathways following CaSR activation were used. The effects of 5 mM [Ca^2+^]_o_ and 10 µM NPS-R568 on the RMP of CHO-Kv7.2/7.3 were investigated before and after the application of each compound. The depolarising effects of 5 mM [Ca^2+^]_o_ and 10 µM NPS-R568 were significantly attenuated by –2.6 ±0.7 mV and –2.3 ±0.8 mV, respectively, in the presence of 100 µM 8-Br-cAMP, a protein kinase A (PKA) activator (Fig 5A and B). In contrast, a non-selective blocker of Ca^2+^ influx NiCl_2_, a chelator of [Ca^2+^]_i_ 1,2-Bis(2-aminophenoxy)ethane-N,N,N’,N’-tetraacetic acid tetrakis acetoxymethyl ester (BAPTA-AM) and a phospholipase C (PLC) inhibitor U73122 had no effect on CaSR-mediated depolarisation (Appendix Fig S2A-D).

**Figure 5.**
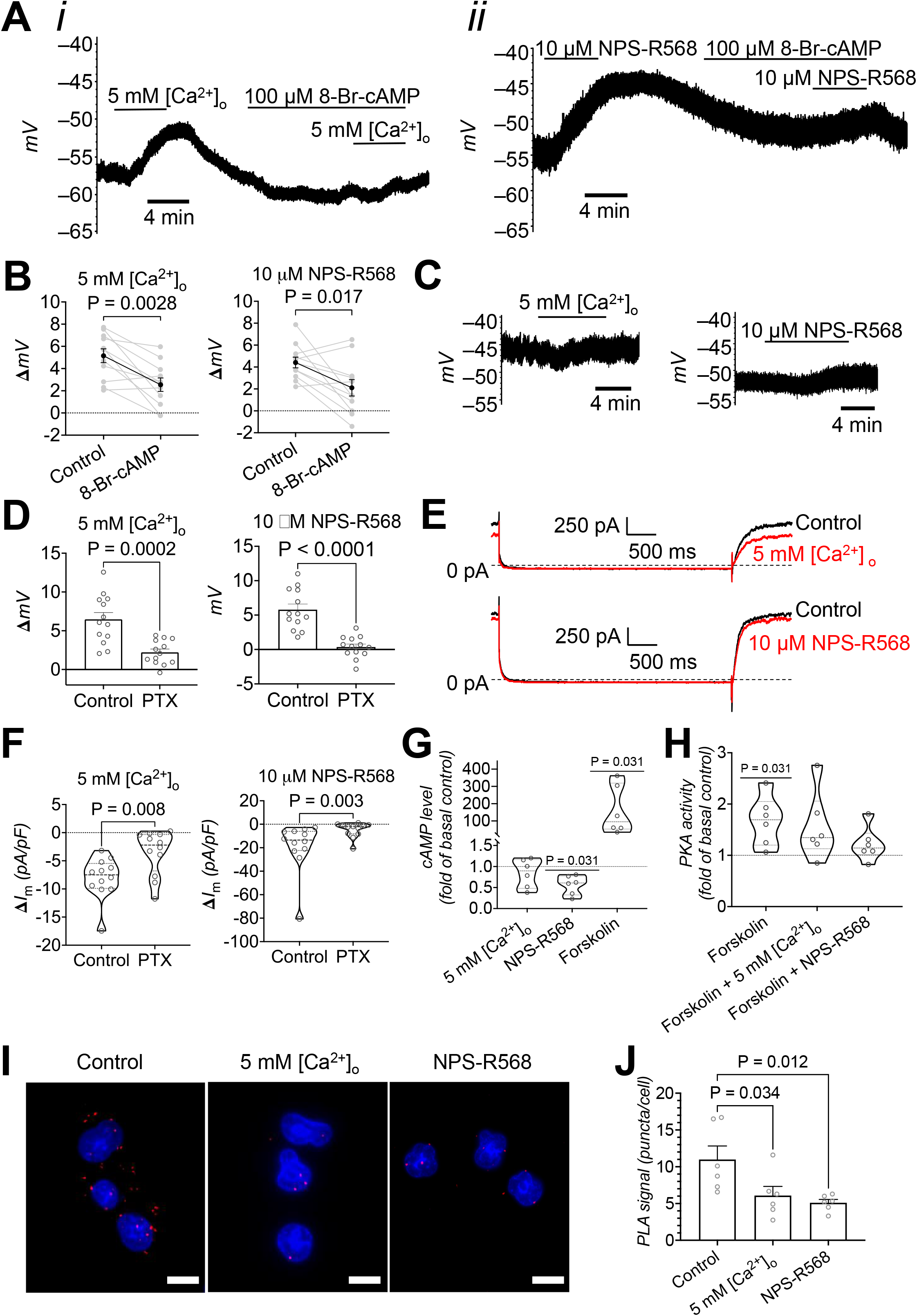
CaSR-Kv7.2/7.3 channel crosslink is mediated through G_i/o_ protein-AC-cAMP-PKA signalling. A Exemplar traces show the effects of 100 μM 8-Br-cAMP on depolarisation by 5 mM [Ca^2+^]_o_ (i) and 10 μM NPS-R568 (ii) in CHO-Kv7.2/7.3. B Graphs compare the magnitude of depolarisation by 5 mM [Ca^2+^]_o_ and 10 μM NPS-R568 before and after the application of 8-Br-cAMP (n=11 cells) in CHO-Kv7.2/7.3. C Exemplar traces show the effects of 5 mM [Ca^2+^]_o_ and 10 μM NPS-R568 on the RMP of CHO-Kv7.2/7.3 after 20-48 hours of 100 ng/ml PTX incubation. D Graphs compare the magnitude of depolarisation by 5 mM [Ca^2+^]_o_ and 10 μM NPS-R568 between control and PTX-treated CHO-Kv7.2/7.3 (n=13 cells). E Exemplar traces show the *I*_m_ of PTX-treated CHO-Kv7.2/7.3 in the control solution and in the presence of 5 mM [Ca^2+^]_o_ and 10 μM NPS-R568. F Graphs compares the magnitude of *I*_m_ density reduction induced by 5 mM [Ca^2+^]_o_ and 10 μM NPS-R568 between control and PTX-treated CHO-Kv7.2/7.3 (n=12 cells). G Graph shows the fold change of intracellular cAMP levels induced by 5 mM [Ca^2+^]_o_, 10 μM NPS-R568 and 10 μM forskolin in CHO-Kv7.2/7.3 (n= 6 experiments). H Graph shows the fold change of PKA activity induced by 10 μM forskolin, 10 μM forskolin plus 5 mM [Ca^2+^]_o_ and 10 μM forskolin plus 10 μM NPS-R568 in CHO-KV7.2/7.3 (n= 6 experiments). I Immunofluorescence images show phosphorylated serine-*KCNQ2* PLA signals of CHO-Kv7.2/7.3 in the control condition, 5 mM [Ca^2+^]_o_ and 10 μM NPS-R568. J Graph compares the density of PLA signals in CHO-Kv7.2/7.3 between three conditions: the control condition, 5 mM [Ca^2+^]_o_ and 10 μM NPS-R568 (n= 6 experiments). Data information: Data are shown as means±SEM. Statistical analyses were performed using two-tailed paired t-test (B), two-tailed unpaired t-test (D), Mann-Whitney U test (F), Wilcoxon signed rank test to a theoretical mean of 1.0 (G and H), and one-way ANOVA with Dunnett’s multiple comparison test to the control (J). Where P is not provided, no significant difference was observed.

### 2.3 CaSR-mediated depolarisation and *I*_m_ reduction of CHO-Kv7.2/7.3 are sensitive to pertussis toxin

The sensitivity of CaSR-mediated depolarisation to PKA activation suggested the involvement of the G_i/o_ protein pathway. To verify this assumption, the effect of 100 ng/ml pertussis toxin (PTX), a G_i/o_ protein inhibitor, on the effects of high [Ca^2+^]_o_ and NPS-R568 in CHO-Kv7.2/7.3 was investigated. The PTX-treated cells exhibited significantly reduced depolarisation induced by both 5 mM [Ca^2+^]_o_ (mean difference: –4.3 ±1.0 mV) and 10 μM NPS-R568 (mean difference: –5.4 ±0.9 mV) (Fig 5C and D). The effects of 5 mM [Ca^2+^]_o_ and 10 μM NPS-R568 on *I*_m_ density in PTX-treated cells were also significantly attenuated by 4.5 ±1.6 pA/pF and 13.9 ±6.4 pA/pF, respectively (Fig 5E and F).

### 2.4 Activation of the CaSR reduces basal intracellular cAMP levels, forskolin-stimulated PKA activity and serine phosphorylation in CHO-Kv7.2/7.3

Coupling of the CaSR to the G_i/o_ protein pathway was further confirmed by investigating the effects of CaSR activation on intracellular cAMP levels and PKA activity in CHO-Kv7.2/7.3. Application of 10 μM NPS-R568 significantly reduced basal intracellular cAMP levels by 0.44 ±0.1 folds, whereas 10 μM forskolin, a transmembrane adenylate cyclase (AC) activator, increased intracellular cAMP levels by 158.6 ±57.2 folds (Fig 5G). In addition, forskolin modestly but significantly increased PKA activity by 1.7 ±0.2 folds; such significance was not apparent when forskolin was applied in combination with either 5 mM [Ca^2+^]_o_ or NPS-R568 (Fig 5H).

The effect of CaSR activation on serine phosphorylation of the Kv7.2 channel subunit was investigated using the proximity ligation assay (PLA). High [Ca^2+^]_o_ and 10 μM NPS-R568 significantly reduced the density of PLA signals by –4.9 ±1.9 punctae/cell and –5.9 ±1.9 punctae/cell, respectively, in CHO-Kv7.2/7.3 (Fig 5I and J). No PLA signal was observed in negative controls where primary antibodies were omitted.

### 2.5 The algogenic cocktail enhances induced excitability of hiPSC-derived nociceptive-like neurons without affecting spontaneous activity

Functional assessment with patch-clamp electrophysiology demonstrated a spectrum of spontaneous and induced excitability in the hiPSC-derived nociceptive-like neurons (Fig 6A-D). To augment the excitability of the hiPSC-derived nociceptive-like neurons, a cocktail of algogenic mediators (1 μM bradykinin, 1 μM substance P, 10 μM adenosine triphosphate [ATP] and 50 ng/ml tumour necrosis factor [TNF]-alpha) associated with neuropathic pain (Ellis & Bennett, 2013) was administered to the neuronal cultures. Following three to four days of incubation with the algogenic cocktail, most of the hiPSC-derived nociceptive-like neurons showed no spontaneous action potential (sAP), which was similar to the controls (Fig 6B). The proportion of the algogenic cocktail-treated neurons with a complete train of induced action potentials (iAPs), however, was more than twice relative to the control (24.5% vs 11.9%) (Fig 6D). In addition, spike analysis of the first iAP was performed in the hiPSC-derived nociceptive-like neurons (Fig 7A-H). The total spike height of the algogenic cocktail-treated neurons was significantly increased by 15.7 ±5.7 mV, compared to the controls (Fig 7E).

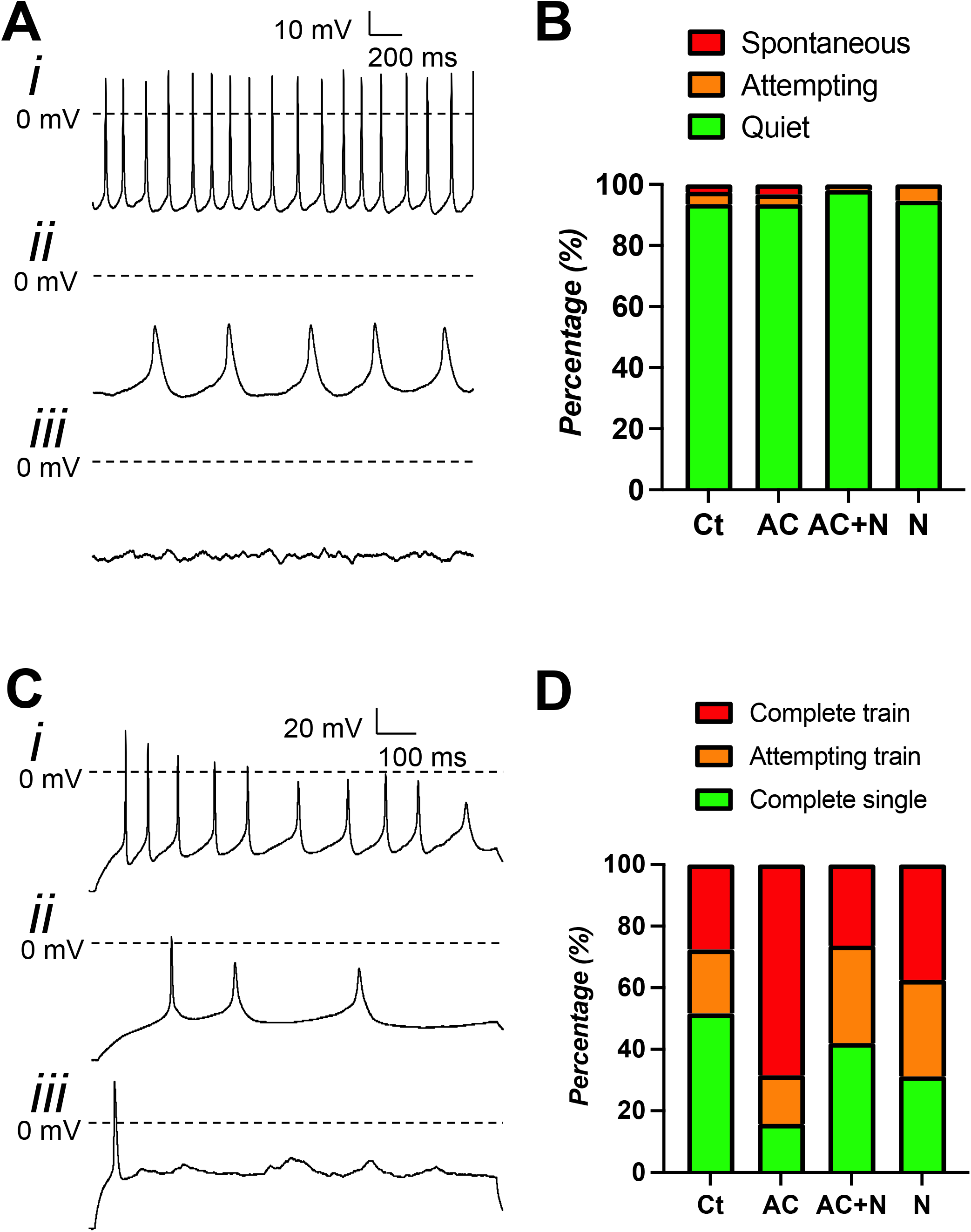

**Figure 7.**
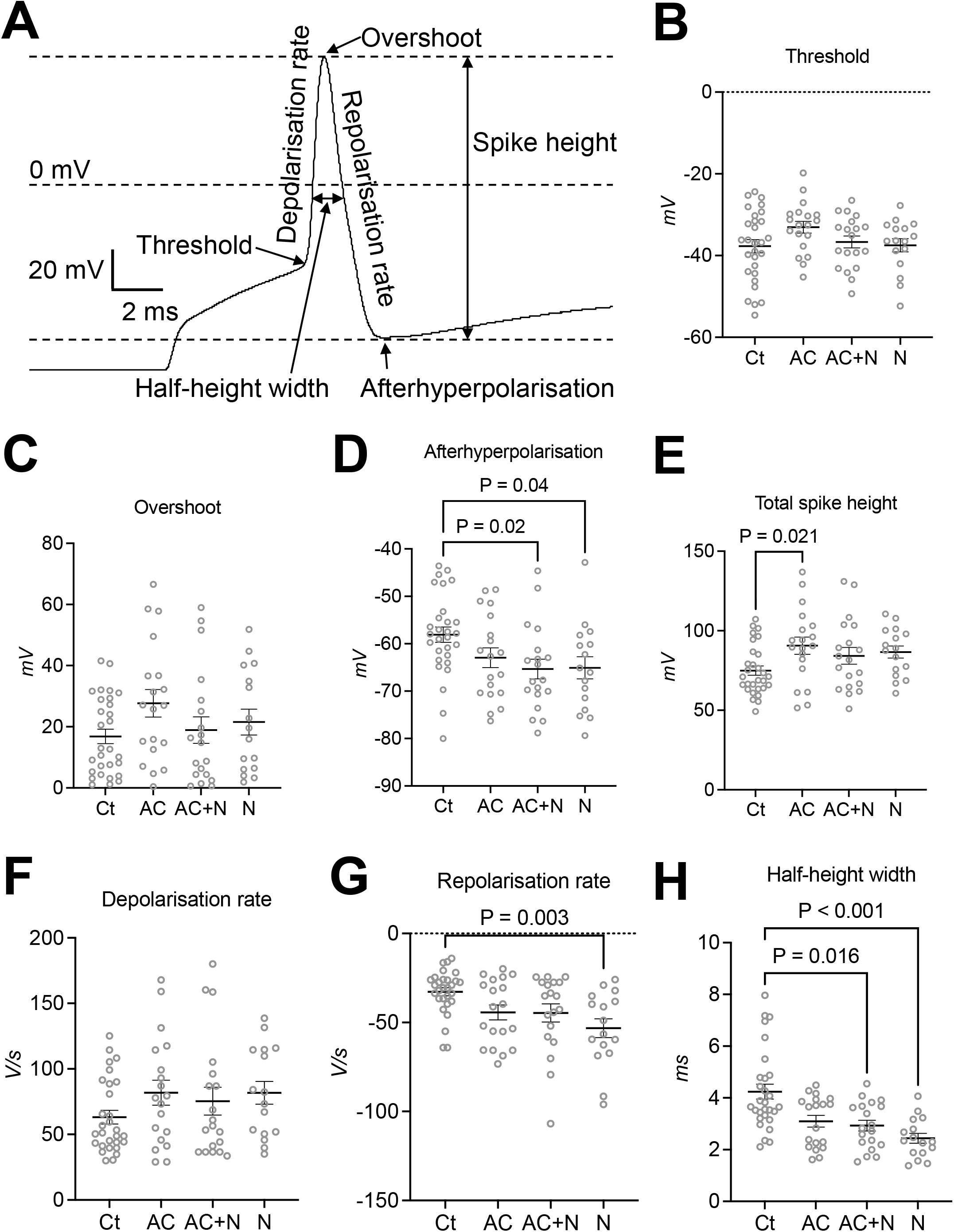
Effects of the algogenic cocktail and NPS-2143 on the characteristics of the first iAP of hiPSC-derived nociceptive-like neurons. A Exemplar trace illustrates the seven characteristics of an iAP investigated: threshold, overshoot, afterhyperpolarisation, total spike height, depolarisation rate, repolarisation rate and half-height width. B-H Graphs compare the seven characteristics of an iAP between four conditions: control (Ct; n=29 cells from seven cultures), the algogenic cocktail (AC; n=19 cells from three cultures), the algogenic cocktail plus NPS-2143 (AC+N; n=19 cells from four cultures) and NPS-2143 (N; n=16 cells from three cultures). Data information: Data are shown as means±SEM. Statistical analyses were performed using one-way ANOVA with Dunnett’s multiple comparison test to the control (B, D, and E) and Kruskal-Wallis test with Dunn’s multiple comparison test to the control (C, F-H). Where P is not provided, no significant difference was observed.

### 2.6 NPS-2143 rescues hiPSC-derived nociceptive-like neurons from hyperexcitability induced by the algogenic cocktail

Like the algogenic cocktail, 1 μM NPS-2143 did not impose a significant effect on the spontaneous excitability of hiPSC-derived nociceptive-like neurons (Fig 6B). When NPS-2143 was applied simultaneously with the algogenic cocktail, a similar distribution of the neurons in each induced excitability classification to the controls was observed, suggesting that NPS-2143 prevented the algogenic cocktail from enhancing the induced excitability (Fig 6D).

Spike analysis of the first iAP showed that NPS-2143 enhanced certain characteristics associated with the repolarisation-hyperpolarisation phase of APs, including afterhyperpolarisation (Fig 7D), repolarisation rate (Fig 7G) and half-height width (Fig 7H). Furthermore, the effect of the algogenic cocktail in enhancing the total spike height was not apparent in the presence of NPS-2143 (Fig 7E).

### 2.7 Algogenic cocktail alters the relationship between the injected current levels and spike frequency in hiPSC-derived nociceptive-like neurons

The relationship between the injected current level and iAP spike frequency (the number of iAPs generated per second) of the hiPSC-derived nociceptive-like neurons exhibiting a complete train of iAPs was analysed in four different conditions (Fig 8A-F). Two-way ANOVA revealed significant effects of both the injected current level and condition on the spike frequency. In the controls, increasing the current amplitude, to a certain extent, correlated with an increase in the spike frequency with two peaks observed at 30 pA and 140 pA (Fig 8A). In the algogenic cocktail-treated neurons, the spike frequency reached its peak at 50 pA then continuously declined with no secondary peak observed (Fig 8B); a similar trajectory was found when the cells were treated in the algogenic cocktail plus NPS-2143 (Fig 8C). The NPS-2143 group showed a similar current-spike frequency relationship to the control, but the two peaks were slightly shifted to 50 pA and 170 pA, respectively (Fig 8D). Post-hoc analysis with Dunnett’s multiple comparison tests revealed the algogenic cocktail reduced the mean iAP spike frequency (mean difference: –2.0 ±0.6 Hz), which was rescued by co-treatment with NPS-2143 (Fig 8F).

**Figure 8.**
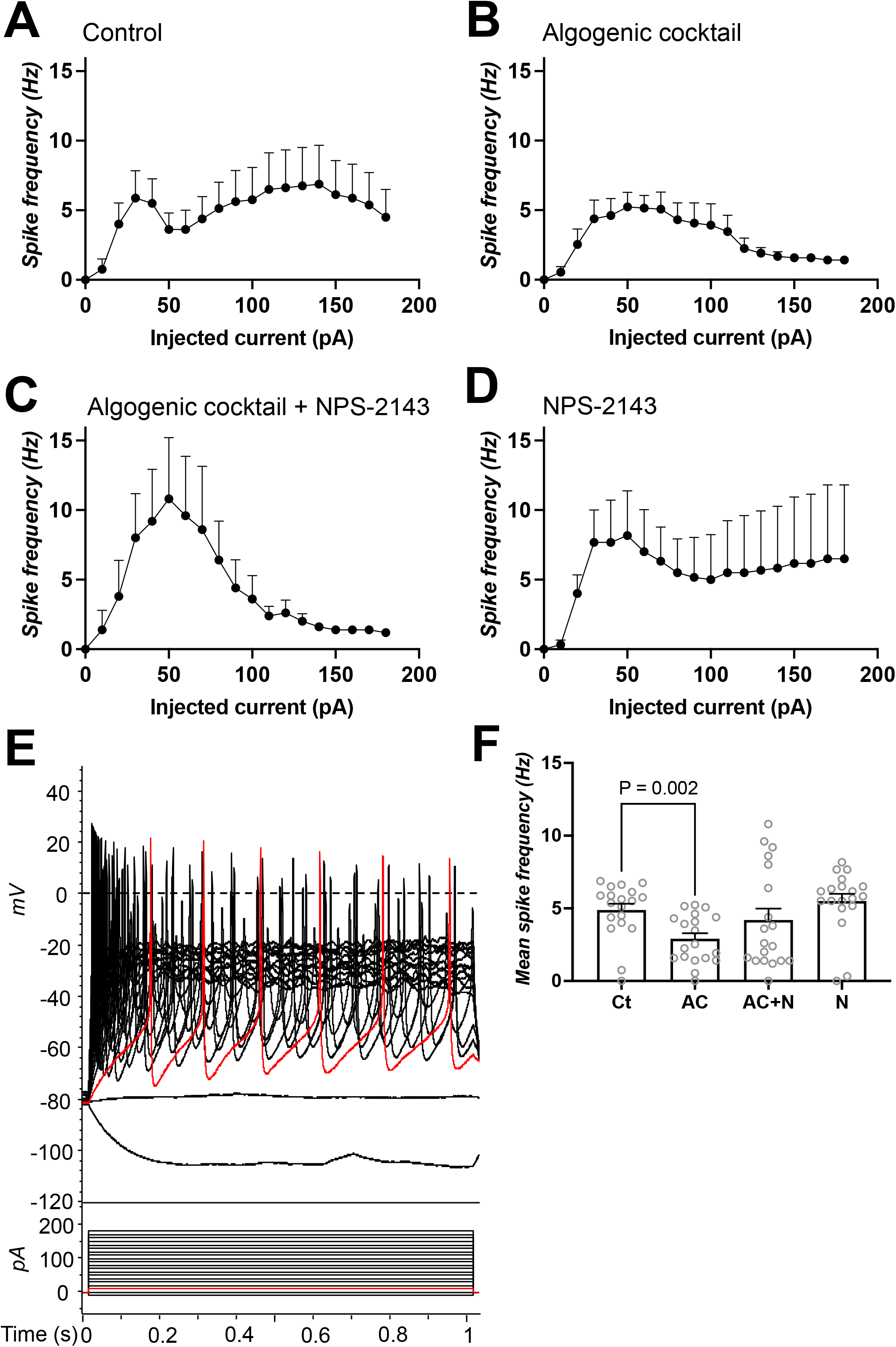
Effects of the algogenic cocktail and NPS-2143 on the iAP spike frequency of hiPSC-derived nociceptive-like neurons. A-D Graphs show the iAP spike frequency at each injected current level of hiPSC-derived nociceptive-like neurons expressing a complete train of iAPs in each condition. E Exemplar traces illustrate trains of induced action potentials in HD33n1 hiPSC-derived nociceptive-like neurons recorded in the current-clamp mode (upper panel) and the 1-s current injection protocol used for recording (lower panel). F Graphs compare the mean iAP spike frequency between four conditions: control (Ct; n=8 cells from five cultures), the algogenic cocktail (AC; n=13 cells from three cultures), the algogenic cocktail plus NPS-2143 (AC+N; n=5 cells from four cultures) and NPS-2143 (N; n=6 cells from three cultures). Data information: Data are shown as means±SEM. Statistical analysis was performed using two-way ANOVA with Dunnett’s multiple comparison test to the control (E). Where P is not provided, no significant difference was observed.

## 3 Discussion

The significance of this study is two-fold: the discovery of the CaSR-Kv7.2/7.3 channel crosslink and the therapeutic potential of targeting the CaSR for neuronal hyperexcitability management. Our conclusions are drawn from experiments involving multiple cell types, which showed congruent results. First, the mechanism by which the CaSR regulates cellular electrical properties via Kv7.2/7.3 channels is originally described in this study. Activation of the CaSR decreased intracellular cAMP levels, forskolin-induced PKA activity and serine phosphorylation of the Kv7.2 channel subunit. Patch-clamp experiments revealed that CaSR activation caused *I*_m_ reduction and depolarisation via inhibition of Kv7.2/7.3 channels. It was demonstrated that CaSR-mediated depolarisation was attenuated by inhibition of G_i/o_ protein or stimulation of PKA. Secondly, the successful differentiation of hiPSCs to nociceptive-like neurons with the ability to produce APs was functionally demonstrated. The algogenic cocktail induced hyperexcitability in hiPSC-derived nociceptive-like neurons as demonstrated by the increased proportion of the neurons exhibiting a complete train of iAPs and enhancement of the iAP total spike height. Such effects of the algogenic cocktail were not apparent when the calcilytic NPS-2143 was present, suggesting that the calcilytic rescued the hiPSC-derived nociceptive-like neurons from algogenic cocktail-induced hyperexcitability.

Our experiments on several cellular models demonstrated that activation of the CaSR by [Ca^2+^]_o_ and NPS-R568 reduced the *I_m_* and caused depolarisation. In the absence of Kv7.2/7.3 channel expression or when the channels were blocked by XE991, CaSR-mediated depolarisation was nearly absent. Collectively, our findings suggest that activation of the CaSR causes depolarisation via Kv7.2/7.3 channel function inhibition. Furthermore, an overlapping functional expression of the CaSR and Kv7 channels observed in a population of mouse DRG neurons in the present study suggested that the CaSR-Kv7 channel crosslink may play a role in regulating neuronal excitability. Since inhibition of neuronal Kv7 channel activity leads to augmented excitability (Liu *et al*., 2010; Peng *et al*., 2017; Yue & Yaari, 2004), our findings fit into the paradigm that activation of the CaSR enhances neuronal excitability (Vyleta & Smith, 2011) and it follows that inhibition of the CaSR with calcilytics may suppress neuronal hyperexcitability.

Our findings are in accordance with previous studies (Chang *et al*., 1998; Di Mise *et al*, 2018) in demonstrating that activation of the CaSR by a PAM agonist NPS-R568 reduced intracellular cAMP levels, confirming the coupling of the CaSR to G_i/o_ protein. High [Ca^2+^]_o_, however, did not have a significant effect in our study. This effect could be explained by the preferential coupling of the CaSR to G_q/11_ protein when the CaSR is stimulated by [Ca^2+^]_o_, compared to NPS-R568 (Cook *et al*, 2015; Nemeth *et al*, 1998). G_q/11_ protein triggers inositol triphosphate-induced [Ca^2+^]_i_ mobilisation, and unlike G_i/o_ protein-sensitive isoforms of transmembrane AC, soluble AC can be activated by risen [Ca^2+^]_i_ and subsequently increase cAMP levels (Tresguerres *et al*, 2011). Therefore, activation of the CaSR can exhibit opposite effects on cAMP levels via G_i/o_ protein and G_q/11_ protein, attenuating the overall response. Given our current understanding that cAMP signalling occurs in discrete intracellular microdomains (Tresguerres *et al*., 2011), the G_i/o_ protein-mediated reduction in cAMP levels near the cell membrane can be masked by an increase in cAMP levels within the cytoplasmic microdomain when measured by enzyme-linked immunosorbent assays (ELISAs).

Since PKA function is dependent on cAMP levels (Søberg & Skålhegg, 2018), CaSR signalling may also interfere with PKA activity. In the present study, stimulation of PKA activity by forskolin was attenuated by NPS-R568 and to a lesser extent by high [Ca^2+^]_o_. Our findings thus confirm the association between CaSR activation and G_i/o_ protein-transmembrane AC-cAMP-PKA signalling. The smaller effect of [Ca^2+^]_o_ was likely pertained to the biased signalling of [Ca^2+^]_o_ towards the G_q/11_ protein pathway (Cook *et al*., 2015; Nemeth *et al*., 1998) and PKA being compartmentalised in a similar fashion to cAMP (Tresguerres *et al*., 2011).

The G_i/o_ protein is sensitive to PTX, which causes adenosine diphosphate ribosylation of the G_i/o_ protein alpha subunit. The ribosylated alpha subunit is then uncoupled from the corresponding GPCR, preventing AC inhibition and subsequent cAMP level reduction (Mangmool & Kurose, 2011). A previous study has shown that the *I_m_*is potentiated by cAMP-dependent PKA phosphorylation at Ser^52^ of the Kv7.2 channel subunit (Schroeder *et al*., 1998). It could then be hypothesised that a reduction in cAMP levels, PKA activity and serine phosphorylation would lead to the inhibition of Kv7.2/7.3 channel activity. In the present study, depolarisation and *I_m_* reduction induced by high [Ca^2+^]_o_ and NPS-R568 were abated in PTX-treated CHO-Kv7.2/7.3. Our data also showed that a PKA activator 8-Br-cAMP lessened CaSR-mediated depolarisation. Furthermore, the PLA showed that activation of the CaSR significantly reduced phosphorylated serine signals associated with the Kv7.2 channel subunit. Altogether, our results suggest that G_i/o_ protein-transmembrane AC-cAMP-PKA signalling pathway mediates the CaSR-Kv7.2/7.3 channel crosslink as illustrated in Fig 9.

**Figure 9.**
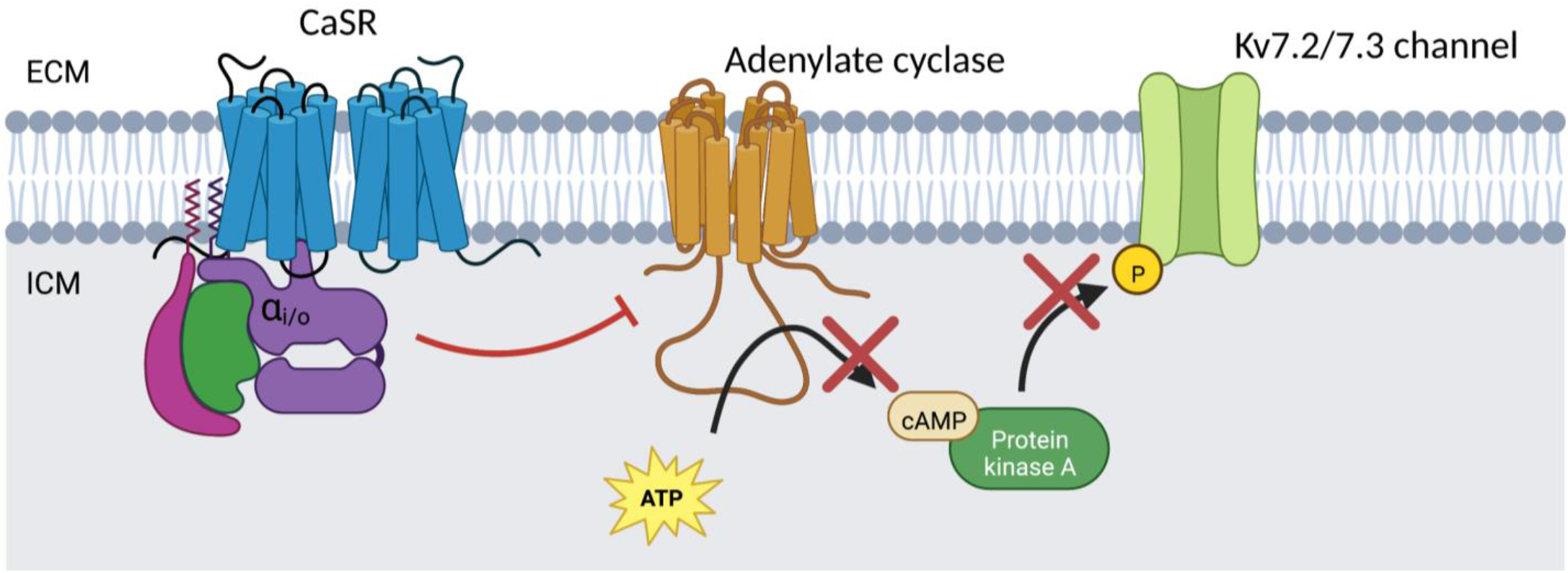
Proposed mechanism of the CaSR-Kv7.2/7.3 channel crosslink. Activation of the CaSR stimulates G_i/o_ protein, which inhibits transmembrane AC. The results are decreases in cAMP levels, PKA activity, serine phosphorylation, and Kv7.2/7.3 channel function as represented by *I_m_* reduction and depolarisation. CaSR, Ca^2+^-sensing receptor; ATP, adenosine triphosphate; cAMP, cyclic adenosine monophosphate; Kv7.2/7.3, voltage-gated K^+^ channel type 7.2/7.3; ECM, extracellular matrix; ICM, intracellular matrix. Created with BioRender.com.

To simulate neuropathic pain *in vitro*, hiPSC-derived nociceptive-like neurons were chronically incubated with the algogenic cocktail to induce hyperexcitability by disrupting Kv7 channel activity as downregulation of neuronal Kv7 channels has been reported in several *in vivo* models of neuropathic pain (Cisneros *et al*, 2015; Laumet *et al*, 2015; Ling *et al*., 2017; Rose *et al*, 2011). The algogenic cocktail thus incorporated the known Kv7 channel-suppressing algogenic mediators, including bradykinin (Liu *et al*., 2010), ATP (Zaika *et al*., 2007) and substance P (Linley *et al*, 2012). In addition, TNF-alpha, a cytokine associated with neuropathic pain, was included (Bowen *et al*, 2006; Koizumi *et al*, 2021). Our data demonstrated that the algogenic cocktail induced hyperexcitability in hiPSC-derived nociceptive-like neurons. Firstly, a higher percentage of the neurons with a complete train of iAPs was observed following algogenic cocktail treatment, suggesting the suppression of the Kv7.2/7.3 channel activity. Our findings are supported by previous studies demonstrating that the iAP spike frequency is enhanced in hiPSC-derived nociceptive-like neurons from patients with inherited erythromelalgia, a persistent pain condition associated with gain-of-function mutations of Nav1.7 channels (Cao *et al*, 2016; Meents *et al*, 2019; Mis *et al*, 2019). Bradykinin, which was a part of our algogenic cocktail, was also shown to enhance iAP firing in isolated human DRG neurons (Davidson *et al*, 2014). Though the precise mechanisms governing the enhancement of the iAP spike frequency were not investigated in this study, it could be speculated that potentiation of Nav1.7 channels might be one of the mechanisms. Secondly, a significant increase in the total iAP spike height was observed following algogenic cocktail treatment, suggesting that the Nav1.7 and Nav1.8 currents were enhanced (Meents *et al*., 2019; Renganathan *et al*, 2001; Savio-Galimberti *et al*, 2014). Unlike the controls, a steady decrease in the iAP spike frequency at high injected current levels was found in the algogenic cocktail-treated hiPSC-derived nociceptive neuron-like cells. A previous study showed that a more hyperpolarised Nav current availability was associated with lower AP firing rates in hiPSC-derived neurons (Telezhkin *et al*, 2016). Hence, it can be hypothesised that the algogenic cocktail induced a hyperpolarising shift of the Nav current availability window, which underlay the observed reduction in the iAP spike frequency at high injected current levels.

Our initial findings about the CaSR-Kv7.2/7.3 channel crosslink posited that in neuropathic pain where the activity of Kv7.2/7.3 channels was suppressed (Cisneros *et al*., 2015; Laumet *et al*., 2015; Ling *et al*., 2017; Rose *et al*., 2011), a calcilytic might help alleviate such circumstance by withdrawing CaSR-mediated inhibition of Kv7.2/7.3 channels and thus restore normal excitability. In our experiments, the effect of the algogenic cocktail in increasing the percentage of hiPSC-derived nociceptive neuron-like cells with a complete train of iAPs was not observed when NPS-2143 was also applied, suggesting that NPS-2143 stabilised neuronal excitability and rescued the neurons from the algogenic cocktail-induced enhancement of iAP firing. This finding further supports our hypothesis that calcilytics may reduce neuronal excitability via the CaSR-Kv7.2/7.3 channel crosslink as Kv7.2/7.3 channel suppression was shown to enhance iAP firing (Gu *et al*, 2005; Peng *et al*., 2017).

Spike analysis of the first iAP revealed that NPS-2143 enhanced afterhyperpolarisation in the present study. Afterhyperpolarisation is mediated by Kv7.2/7.3 channels and small/intermediate-conductance Ca^2+^-activated K^+^ channels (Gu *et al*., 2005; Mateos-Aparicio *et al*, 2014; Savić *et al*, 2001; Storm, 1989; Tzingounis & Nicoll, 2008). However, Vassilev *et al*. (1997) demonstrated that activation of the CaSR increased the open-state probability of Ca^2+^-activated K^+^ channels. Therefore, NPS-2143 likely enhanced afterhyperpolarisation via Kv7.2/7.3 channels rather than Ca^2+^-activated K^+^ channels, further supporting our theory of the CaSR-Kv7.2/7.3 channel crosslink. Furthermore, NPS-2143 enhanced the repolarisation rate, suggesting that calcilytics via the CaSR may mediate the function of AP repolarisation-regulating ion channels such as Kv2 and Kv3 channels (Johnston *et al*, 2010; Tsantoulas & McMahon, 2014). Regulation of Kv3 channels via CaSR signalling was previously demonstrated in neurons (Kaczmarek & Zhang, 2017; Ritter *et al*, 2012). Thus, our findings provide new insight for how the CaSR might be an appropriate target for neuropathic pain management.

Though NPS-2143 inhibited the algogenic cocktail in driving the hiPSC-derived nociceptive-like neurons to develop a complete iAP train characteristic, it appeared to synergise with the algogenic cocktail in increasing the iAP spike frequency at lower current injection of the neurons that presumably already expressed a train of iAPs. Blocking of Ca^2+^-activated K^+^ channels was shown to enhance iAP firing in rodent DRG neurons (Pagadala *et al*, 2013; Tsantoulas & McMahon, 2014; Zhang *et al*, 2003). Based on existing evidence that stimulation of the CaSR upregulates Ca^2+^-activated K^+^ channel function (Vassilev *et al*., 1997), one might speculate that NPS-2143, via the CaSR, inhibits Ca^2+^-activated K^+^ channels, leading to augmented iAP firing. A previous study demonstrated that most human DRG neurons (not quantified) produced a single iAP (Davidson *et al*., 2014). Therefore, when it comes to modulation of collective excitability, the effect of NPS-2143 in preventing algogenic cocktail-induced transformation of the neurons with a single iAP towards multiple iAPs may be greater than its effect in enhancing iAP firing of the cells already at the high-end of the excitability spectrum. The effect of the calcilytic on neuronal excitability, thus, is likely to be multimodal with several knowledge gaps worth pursuing in the future.

Some limitations can be recognised in this study. First, the observed effect of [Ca^2+^]_o_ could be due to its off-target actions beyond the CaSR. This limitation was circumvented by using a specific PAM agonist of the CaSR, NPS-R568, to supplement our findings and the same effects of both [Ca^2+^]_o_ and NPS-R568 were observed. The effect of [Ca^2+^]_o_, however, is relevant and still deserves attention as [Ca^2+^]_o_ is the dominant natural agonist of the CaSR in the body. In the event of hypercalcemia or inflammation where [Ca^2+^]_o_ substantially rises (Kaslick *et al*, 1970; Lansdown *et al*, 1999), its stimulatory effect on the CaSR could have a significant impact on neuronal excitability. Fluctuations of [Ca^2+^]_o_ can also modulate synaptic transmission (Phillips *et al*, 2008), signifying the importance of [Ca^2+^]_o_. Secondly, ELISAs were used to quantify intracellular cAMP levels and PKA activity. This technique only gives information about overall changes within the cell but no information regarding localisation of the effects. This warrants future investigations into the effect of CaSR activation on spatially distinct cAMP levels and PKA activity. The hiPSC-derived nociceptive neuron-like model provides a promising platform to explore novel therapeutics for pain but is limited by how well the model can replicate *in vivo* human nociception. Our model still lacked the full complexity of *in vivo* models including: the interaction with other cell types such as glia, which were shown to play a crucial role in the modulation of neuronal excitability and neuropathic pain development (Zhang *et al*, 2017), the complete assemblage of algogenic mediators released *in vivo* (Ellis & Bennett, 2013) and the complexity of the biopsychosocial model of pain (Leeuw & Klasser, 2018).

To summarise, this study demonstrates the CaSR-Kv7.2/7.3 channel crosslink mechanism and the calcilytic’s potential for controlling augmented excitability in the hiPSC-derived nociceptive-like neuronal model. Our findings spearhead the CaSR as a potential therapeutic target for Kv7 channel-associated hyperexcitability disorders such as neuropathic pain, which is worth further investigation.

## 4 Materials and methods

### 4.1 Compounds and antibodies

The following compounds and antibodies were used in this study: NPS-R568, a CaSR PAM agonist (Tocris Bioscience, Abingdon, UK); NPS-2143, a CaSR NAM (Tocris); retigabine, a Kv7 channel opener (Tocris); XE991, a Kv7 channel blocker (Tocris); forskolin, a transmembrane AC activator (Tocris); 8-Br-cAMP, a phosphodiesterase-resistant cAMP analogue and PKA activator (Merck Life Science, Dorset, UK); *Bordetella* PTX, a specific inhibitor targeting the alpha subunit of the G_i/o_ protein (Merck); U73122, a PLC inhibitor (Tocris); and BAPTA-AM, an [Ca^2+^]_i_ chelator (Tocris); NiCl_2_ hexahydrate, a non-selective blocker of Ca^2+^ influx (Fisher Scientific, Loughborough, UK); bradykinin (Merck); substance P (Tocris); ATP (Merck); TNF-alpha (Merck); mouse monoclonal anti-CaSR (ab19347, 1:100, Abcam, Cambridge, UK); rabbit polyclonal anti-TRPV1 (ab3487, 1:100, Abcam); rabbit polyclonal anti-*KCNQ2* (ab22897, 1:200, Abcam); rabbit polyclonal anti-*KCNQ3* (PA1-930, 1:100, Life Technologies, Paisley, UK); Alexa 488 donkey anti-mouse IgG (H+L) highly crossed-adsorbed secondary antibody (A21202, 1:200, Life Technologies); Alexa 594 donkey anti-rabbit IgG (H+L) highly crossed adsorbed secondary antibody (A21207, 1:200, Life Technologies); rabbit polyclonal anti-CaSR (SAB4503369, 1:100, Merck); mouse monoclonal anti-MAP2 (M1406, 1:100, Merck); Alexa 488 goat anti-mouse IgG (H+L) pre-adsorbed secondary antibody (ab150117, 1:250, Abcam); rabbit polyclonal anti-*KCNQ2* (ab22897, 1:200, Abcam); mouse monoclonal anti-phosphoserine conjugated to Duolink® PLA MINUS probe (DUO87012, 1:50, Merck).

### 4.2 Cell line culture

CHO-Kv7.2/7.3 and HEK-CaSR were cultured separately in Gibco™ Dulbecco’s Modified Eagle Medium (DMEM)/Nutrient Mixture F-12 (Life Technologies) supplemented with 10% foetal bovine serum (Life Technologies), 1% L-glutamine (Life Technologies), and 1% penicillin/streptomycin (Life Technologies) and maintained in a humidified incubator at 37°C with 5% CO_2_/95% O_2_. The culture medium was replaced every two to three days and confluency was assessed with a Leica DM IL inverted microscope (Leitz, Wetzlar, Germany). The cells were passaged using 0.25% trypsin-ethylenediamine tetra acetic acid (EDTA; Life Technologies) upon reaching 80% confluency.

HD33n1 hiPSCs were cultured in 6-well plates, which were precoated with Corning® Matrigel® basement membrane matrix (Scientific Laboratory Supplies, Nottingham, UK), containing mTeSR™ plus medium (Stemcell technologies, Cambridge, UK) at 37 °C with 5% CO_2_/95% O_2_. The culture medium was replaced every two to three days; the cells were passaged using 0.02% EDTA Ca^2+^/Mg^2+^-free PBS pH 7.2-7.4 (Merck) once 80% confluency was reached. The hiPSCs were differentiated into neurons using a previously published protocol (Telezhkin *et al*., 2016) with minor modifications: incorporation of 0.33 μl/ml ECMAtrix-511 E8 laminin substrate (Merck) into the SCM2 medium and exclusion of LM22A4 from both SCM1 and SCM2 media. After at least 10 days in SCM2 medium, the cells were used for experiments.

### 4.3 Isolation and culture of primary DRG neurons

Animal procedures were approved by the Animal Welfare and Ethical Review Body of Newcastle University. Naïve 6-month-old male CB57BL/6 mice (n=6 mice, Comparative Biology Centre, Newcastle University, UK) were euthanised by cervical dislocation and their DRG extracted from all spinal levels under a microscope. The collected DRG were enzymatically digested using collagenase (5 mg/ml, Merck), papain (5mg/ml, Merck), and dispase (10 mg/ml, Merck) diluted in Ca^2+^/Mg^2+^-free Hanks’ balanced salt solution (Life Technologies) for 40 minutes (min) at 37°C. The DRG were gently triturated and resuspended in DMEM/Nutrient Mixture F-12. The DRG cell suspension was plated onto circular glass coverslips (Fisher Scientific), which were pre-coated sequentially with poly-D-lysine & laminin (Merck) and Matrigel, and fitted into 24-well plates. The isolated DRG neurons were cultured *in vitro* in 50% DMEM/Nutrient Mixture F-12 and 50% Neurobasal-A medium (Life Technologies), supplemented with 1% L-glutamine, 1% penicillin/streptomycin, 2% MACS® NeuroBrew-21 (Miltenyi Biotec, Surrey, UK), 2 μM PD0332991 (Tocris), 10 ng/ml human BDNF (Miltenyi Biotec), 3 μM CHIR-99021 (Tocris), 200 μM L-ascorbic acid (Merck), and CaCl_2_ (to the final concentration of 1.8 mM). Experiments were carried out three to 14 days after plating.

### 4.4 Immunocytochemistry

The cells (CHO-Kv7.2/7.3, HEK-CaSR, or hiPSC-derived neurons) were plated on 13 mm-diameter circular glass coverslips, fixed with 4% paraformaldehyde (Fisher Scientific) for 15 min and washed with phosphate buffer solution (PBS) three times. The fixed cells were blocked and permeabilised with 1% bovine serum albumin (Merck) and 0.1% Triton™ X-100 (Merck) in PBS for one hour at room temperature. The coverslips were then incubated with the primary antibody diluted in the blocking/permeabilisation solution at a concentration as indicated above and kept at 4 °C overnight. The next day, the coverslips were washed with PBS three times followed by an application of the secondary antibody for one hour at room temperature. Afterwards, the coverslips were washed with PBS three times before being mounted using ProLong™ Diamond Antifade Mountant with 4′,6-diamidino-2-phenylindole (DAPI; Life Technologies). Images were obtained with Olympus BX61 fluorescence microscope or Zeiss LSM800 laser confocal scanning microscope and processed using cellSens imaging software (Olympus, Tokyo, Japan).

### 4.5 Electrophysiology

Prior to electrophysiological experiments, specialised silicone chambers secured on square glass coverslips (approximate volume of 200 μl) were prepared. CHO-Kv7.2/7.3 or HEK-CaSR were then plated on the prepared chambers and incubated at 37°C with 5% CO_2_/95% O_2_ overnight before conducting any electrophysiological experiments. Circular glass coverslips with mouse DRG neurons cultured *in vitro* or hiPSC-derived nociceptive-like neurons were placed inside the silicone chambers prior to conducting experiments.

The chambers were mounted on a microscope stage and perfused with standard bath solution containing (in mM): 146 NaCl 2.5 KCl, 0.5 MgCl_2_, 0.5 CaCl_2_, 10 D-glucose, 5 HEPES; pH was adjusted to 7.4 using NaOH. The cells were patched with borosilicate glass pipettes (Harvard apparatus, Cambourne, UK), with 3 to 7 MΩ resistance when filled, in the whole-cell configuration at room temperature. The standard pipette solution contained (in mM): 140 KCl, 6 NaCl, 4 Na_2_-ATP, 3 MgCl_2_, 1 CaCl_2_, 5 HEPES and 5 EGTA; pH was adjusted to 7.2 with KOH. All salts and compounds in the standard bath and pipette solutions were purchased from Fisher Scientific.

The RMP was recorded with the gap free protocol (current = 0 pA) in a current-clamp mode using Axopatch 200A amplifier and Digidata 1440 A/D interface (Axon Instruments, Forster City, USA). The macroscopic transmembrane *I*_m_ was recorded with the voltage-stepped deactivation protocol by holding the cells at a potential of –20 mV and stepped down to –60 mV for 4 s. All recordings were filtered at 2 kHz and digitised at 5 kHz. Clampex 10.2 (Molecular Devices, Sunnyvale, USA) was used for data acquisition.

In the voltage clamp mode, macroscopic Nav and Kv currents were recorded using a standard voltage-step activation/inactivation protocol as shown in Appendix Fig S1A. The cells were held at –70 mV followed by a 200-ms step starting at –120 mV and increasing by 5 mV per sweep up to 80 mV. After the first activation step, the cells were stepped to 0 mV for 200 ms. The cells were then shifted back to –70 mV for 100 ms before starting the next sweep. Activation of the Nav current was observed as a fast inwards spike at the beginning of the activation step from –70 mV; the magnitude of the activating Nav current was measured at the trough of the inwards spike. Similarly, the inactivating Nav current was measured at the trough of the second inwards spike when the cells were stepped to 0 mV (Appendix Fig S1A). Conductance (*G*) was calculated as the ratio of the current to the driving force; the driving force of the Nav current was calculated by subtracting its equilibrium potential (66.68 mV) from the command voltage. Measurements of Kv currents were taken by averaging the steady state currents during the last 50 ms of the activation voltage step.

For the determination of the spontaneous activity, the cells were held with a current of 0 pA for 1 min to allow for observation of any sAP being generated. The cells were categorised into three groups based on the sAP type: quiet, attempting sAP and complete sAP (Fig 6A) (Telezhkin *et al*., 2016). The ability of the cells to generate iAPs was determined using a 1-s current step injection protocol. First, the cells were artificially hyperpolarised and held between –80 and –90 mV. Afterwards, the current step was incrementally increased by 10 pA per sweep up to a maximum of 180 pA; 30 ms was allowed between steps for recovery. The change in the RMP was observed and the cells were categorised into three groups: complete single iAP, attempting train of iAP and complete train of iAP (Fig 6C) (Telezhkin *et al*., 2016). The cells expressing immature iAPs that did not reach 0 mV were excluded from analyses.

Spike analysis of the first iAP was performed on the cells that expressed at least a complete single iAP. Using Clampfit 10.2, several characteristics of an iAP were determined as shown in Fig 7A. Some characteristics can be measured directly, including overshoot, afterhyperpolarisation and spike height. To determine the half-height width, the voltage at 50% of the spike height was first measured, and the time difference between which this voltage was reached during depolarisation and repolarisation was calculated. The depolarisation rate and repolarisation rate were defined as the maximum rising slope and maximum declining slope, respectively, which were measured from the first differentiation of voltage with respect to time. The threshold was determined at the peak of the third differentiation of voltage with respect to time during the depolarisation phase of an AP (Henze & Buzsáki, 2001; Telezhkin *et al*., 2016).

### 4.6 Detection of intracellular cAMP levels and PKA activity

CHO-Kv7.2/7.3 were plated in 12-well plates at a density of 5×10^5^ cells per well and incubated at 37°C with 5% CO_2_/95% O_2_. The following day, the wells were divided into groups and received appropriate media (Fig 5G and H). The following commercial kits were used to determine intracellular cAMP levels and PKA activity, respectively, according to the manufacturers’ instructions: Enzo® Life Sciences Direct cAMP ELISA kit (ADI-900-066, Fisher Scientific) and Invitrogen™ PKA Colorimetric Activity kit (EIAPKA, Life Technologies). The intracellular cAMP levels and PKA activity were normalised to total protein concentrations, which were determined using the Pierce™ BCA Protein Assay Kit (Life Technologies) for samples prepared for cAMP detection and the Pierce™ 660 nm Protein Assay Reagent (Life Technologies) for samples prepared for PKA activity detection according to the manufacturers’ instructions. Each assay was performed in duplicate.

### 4.7 Phosphoserine-*KCNQ2* PLA

CHO-Kv7.2/7.3 were plated on 13 mm-diameter circular glass coverslips fitted in 24-well plates at a density of 2×10^4^ cells per coverslip and incubated at 37 °C with 5% CO_2_/95% O_2_. The following day, the cells were treated with either 5 mM [Ca^2+^]_o_ or 10 µM NPS-R568 for 4 hours, fixed with 4% paraformaldehyde for 15 min, and stored in 100% methanol at –20 °C. A commercial Duolink® PLA system (Merck) was used according to the manufacturer’s protocol. The following antibodies were used: rabbit polyclonal anti-*KCNQ2* (ab22897, 1:200) and mouse monoclonal anti-phosphoserine conjugated to Duolink® PLA MINUS probe (DUO87012, 1:50). Olympus BX61 fluorescence microscope was used to capture three random areas per coverslip; the density of PLA signals (red fluorescent puncta/cell) were counted using ImageJ software.

### 4.8 Transient transfection of HEK-CaSR with *KCNQ2/3* cDNA

HEK-CaSR were plated on specialised silicone chambers as mentioned above and incubated at 37°C with 5% CO_2_/95% O_2_ overnight. Transfection was performed the following day. In an Eppendorf tube, 2 μl of *KCNQ2/3* cDNA plasmid (0.9 g/l) and 1 μl of pmax green fluorescent protein vector (0.5 g/l) (Lonza, Cologne, Germany) were diluted in 125 μl of Opti-MEM medium (Life Technologies). In a separate tube, 5 μl of Lipofectamine™ 2000 transfection reagent (Life Technologies) was diluted in 125 μl of Opti-MEM® medium and incubated at room temperature for 5 min. The diluted cDNA and diluted Lipofectamine^TM^ 2000 transfection reagent were gently mixed then incubated at room temperature for a further 20 min. Approximately 35 μl of the mixture was added to each chamber and all chambers were incubated at 37°C with 5% CO_2_/95% O_2_. The culture medium in each chamber was then replaced the next day. For the control group, *KCNQ2/3* cDNA was omitted.

### 4.9 Chronic incubation of hiPSC-derived nociceptive-like neurons with the algogenic cocktail and/or the calcilytic NPS-2143

To induce hyperexcitability of hiPSC-derived nociceptive-like neurons, a cocktail of algogenic mediators (1 μM bradykinin, 1 μM substance P, 10 μM ATP and 50 ng/ml TNF-alpha) diluted in SCM2 medium was applied in culture for three to four days. To investigate the effect of a calcilytic on excitability, 1 μM NPS-2143 was applied both alone and in combination with the algogenic cocktail to separate cultures of hiPSC-derived neuron-like cells for three to four days. Analyses of the spontaneous activity, iAP type, characteristics of the first iAP and spike frequency were performed.

### 4.10 Data and statistical analyses

All data are presented as mean ± standard error of the mean (SEM). Data were analysed and visualised using Clampfit 10.2 (Molecular Devices) and GraphPad Prism 9 (GraphPad Software, San Diego, USA).

For the analysis of concentration-response experiments, the voltage change induced by different concentration of [Ca^2+^]_o_ relative to 0.1 mM [Ca^2+^]_o_ was fitted with the Hill equation:

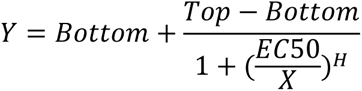

where *Y* is the voltage change, *X* is the [Ca^2+^]_o_, EC_50_ is the half maximal depolarisation response, and *H* is Hill slope.

For activation and inactivation curves, *G* was normalised to the maximum conductance (*G_max_*) and the curves were generated by plotting the normalised conductance (*G/G_max_*) against voltage. Using Microcal Origin (OriginLab, Massachusetts, USA), the conductance-voltage relationship was fitted with the Boltzmann equation:

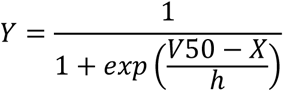

where *Y* is the normalised conductance, *X* is the command voltage, *V50* is the voltage pertaining to the half value of the maximum conductance, and *h* is the slope factor.

Statistical tests (one-sample t-test, Wilcoxon signed rank test, paired t-test, unpaired t-test, Mann-Whitney U test, one-way ANOVA, two-way ANOVA and Kruskal-Wallis test as stated in text and figure legends) were carried out in GraphPad Prism 9; statistical significance differences were recognised at *P*<0.05.

## 5 Acknowledgements

We thank Prof. David A. Brown, UCL for sharing CHO-Kv7.2/7.3 and the *KCNQ2/3* cDNA plasmid, Prof. Daniela Riccardi, Cardiff University for sharing HEK-CaSR, and both for helpful discussions of the results. We thank Prof. Nicholas D. Allen, Cardiff University for sharing the HD33n1 hiPSCs. We would like to acknowledge Chloe Sorrell, Newcastle University for helping with immunocytochemistry, Dr Junyong Huang, Newcastle University for his technical advice on the PLA and Dr Kim Pearce, Newcastle University for her statistical support. This study was supported by a PhD studentship to N.C. from the Anandamahidol Foundation, Thailand, and Wellcome Trust [221678/Z/20/Z] to P.Y. This project was supported in part by Chinese Academy of Sciences President’s International Fellowship Initiative to P.Y. [2021VBC0009] and V.T. [2022VBB0002].

## 6 Author contributions

**Nontawat Chuinsiri**: conceptualisation; data curation; formal analysis; funding acquisition; investigation; methodology; visualisation; writing – original draft; writing – review and editing. **Nannapat Siraboriphantakul**: investigation. **Luke Kendall**: investigation. **Polina Yarova**: conceptualisation; funding acquisition; investigation; methodology; resources; writing – review and editing. **Christopher J. Nile**: conceptualisation; methodology; supervision; writing – review and editing. **Bing Song**: writing – review and editing. **Ilona Obara**: resources; writing – review and editing. **Justin Durham**: conceptualisation; funding acquisition; methodology; supervision; writing – review and editing. **Vsevolod Telezhkin**: conceptualisation; data curation; formal analysis; funding acquisition; investigation; methodology; resources; supervision; visualisation; writing – review and editing.

## 7 Disclosure and competing interests statement

The authors declare that they have no conflict of interest.

## 8 Data availability

Data needed to evaluate the conclusions in the paper are present in the paper. Data and materials generated for this study are available from the corresponding author upon reasonable request. This study includes no data deposited in external repositories.

## Supplemental figure legends

**Supplemental Figure S1.**
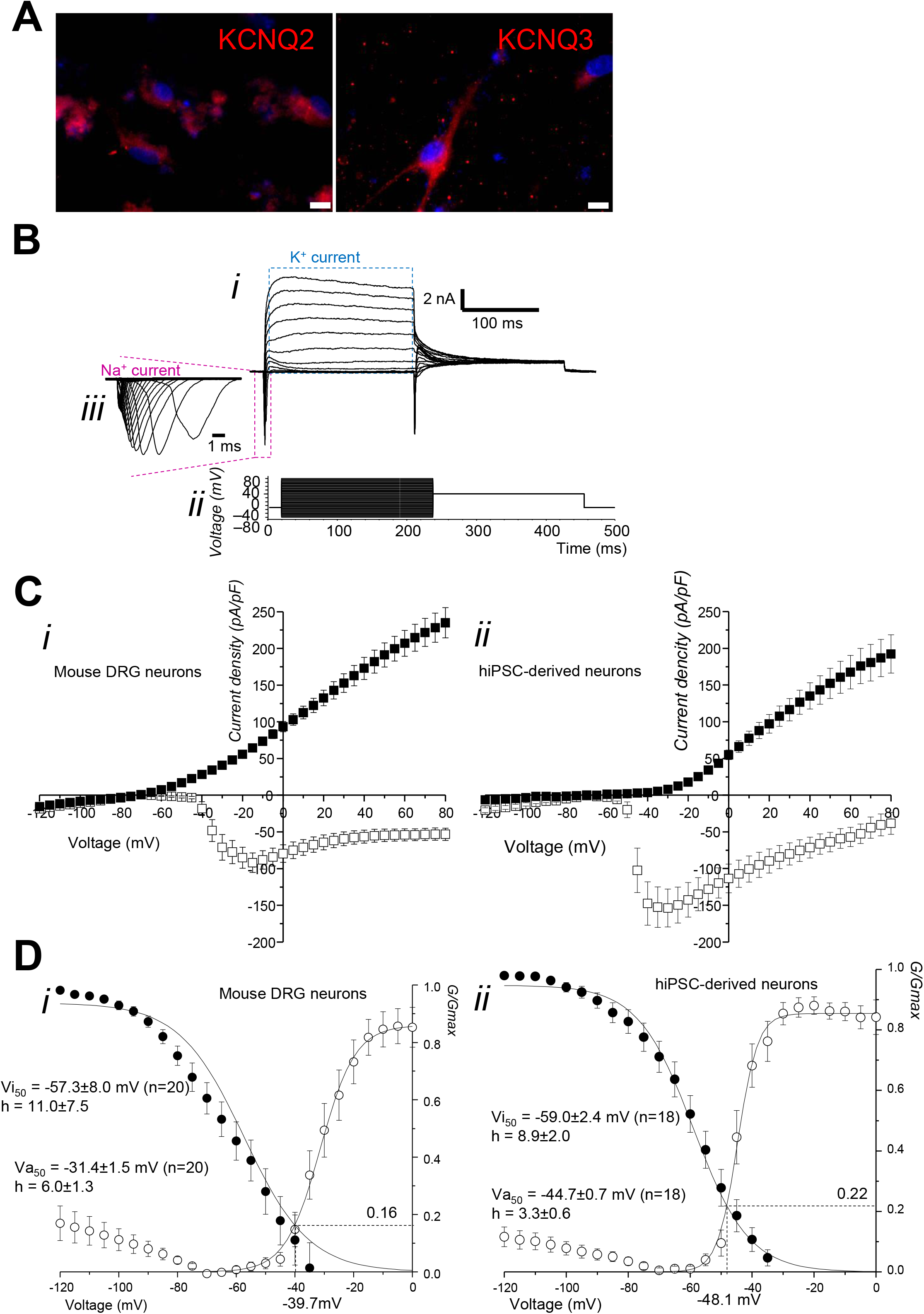
Immunocytochemical and electrophysiological characterisation of mouse DRG neurons and hiPSC-derived neurons. A Immunofluorescence image shows KCNQ2 (ab22897) and KCNQ3 (PA1-930) expressed in hiPSC-derived neurons. The negative control is shown in Fig 4B. Images were obtained with Olympus BX61 fluorescence microscope and are representative of three independent experiments. Scale bars: 10 µm. B Exemplar trace shows macroscopic Nav and Kv currents (i) recorded using the voltage-step activation/inactivation protocol by holding the neurons at –70 mV followed by a 200-ms step starting at –120 mV and increasing by 5 mV per sweep up to 80 mV. After the first activation step, the neurons were stepped to 0 mV for 200 ms. The neurons were then shifted back to – 70 mV for 100 ms before starting the next sweep (ii). A portion of the trace is expanded to show inwards spikes corresponding to the activating Nav current (iii). C Graphs illustrate the mean of maximum Nav (empty squares) and steady-state Kv (filled squares) current density plotted against the voltage of mouse DRG neurons (i) and hiPSC-derived nociceptive-like neurons (ii). D Graphs illustrate the mean of normalised conductance plotted against the voltage for Nav activation (empty circles) and inactivation (filled circles) of mouse DRG neurons (i) and hiPSC-derived nociceptive-like neurons (ii). Data information: Data are shown as means±SEM.

**Supplemental Figure S2.**
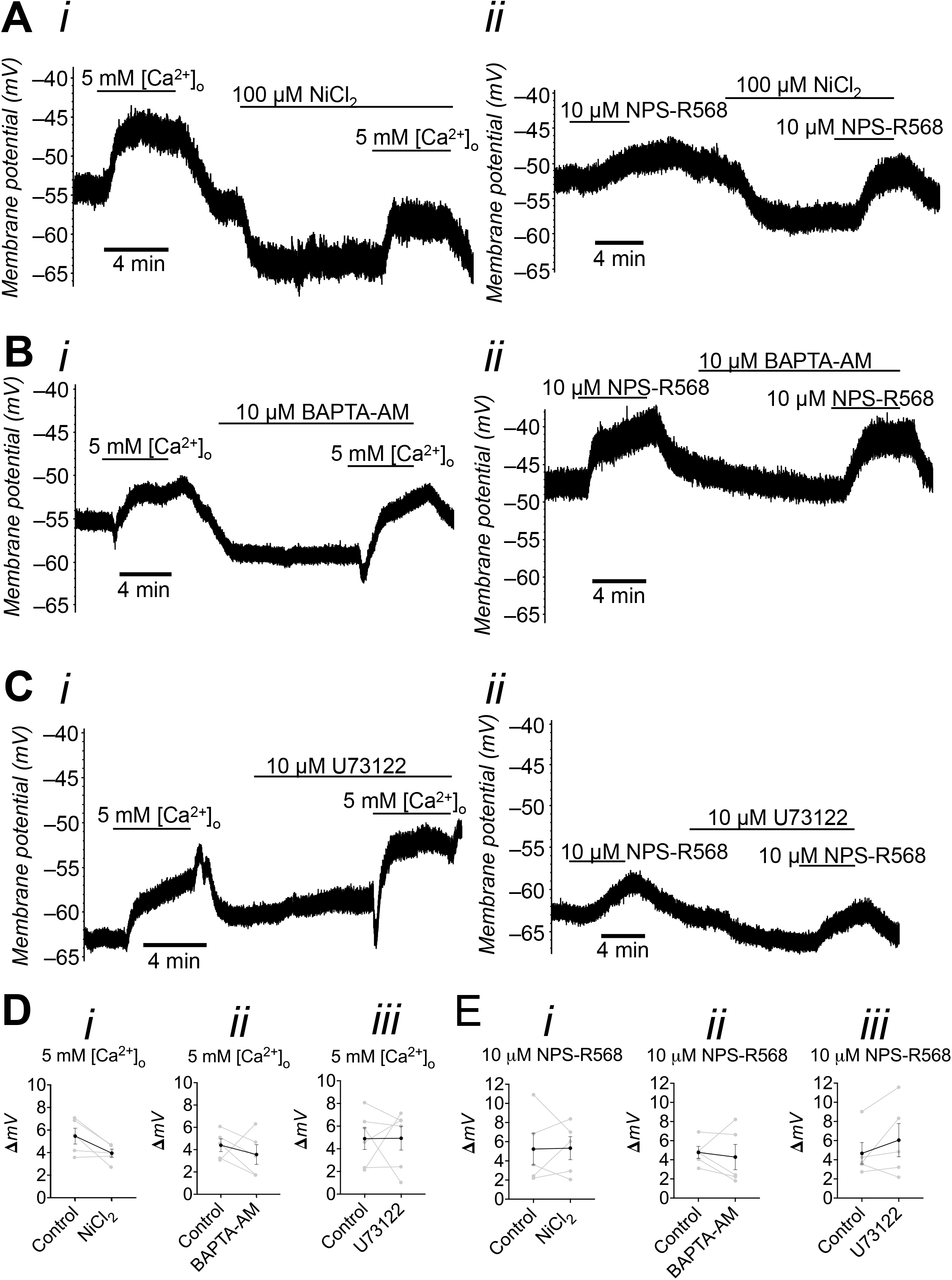
Effects of NiCl_2_, BAPTA-AM and U73122 on CaSR-mediated depolarisation in CHO-Kv7.2/7.3. A Exemplar traces show the effects of 100 μM NiCl_2_ on depolarisation by 5 mM [Ca^2+^]_o_ (i) and 10 μM NPS-R568 (ii) in CHO-Kv7.2/7.3. B Exemplar traces show the effects of 10 μM BAPTA-AM on depolarisation by 5 mM [Ca^2+^]_o_ (i) and 10 μM NPS-R568 (ii) in CHO-Kv7.2/7.3. C Exemplar traces show the effects of 10 μM U73122 on depolarisation by 5 mM [Ca^2+^]_o_ (i) and 10 μM NPS-R568 (ii) in CHO-Kv7.2/7.3. D Graphs show the magnitude of depolarisation by 5 mM [Ca^2+^]_o_ (n=5-6 cells from three to five cultures) before and after the application of each compound (i: NiCl_2_, ii: BAPTA-AM and iii: U73122). E Graphs show the magnitude of depolarisation by 10 μM NPS-R568 (n=5 cells from two to five cultures) before and after the application of each compound (i: NiCl_2_, ii: BAPTA-AM and iii: U73122). Data information: Data are shown as means±SEM. Statistical analyses were performed using two-tailed paired t-test (D and E). Where P is not provided, no significant difference was observed.

